# Inhibition of heme biosynthesis triggers cuproptosis in acute myeloid leukaemia

**DOI:** 10.1101/2024.08.11.607520

**Authors:** Alexander C. Lewis, Emily Gruber, Rheana Franich, Jessica Armstrong, Madison J. Kelly, Carlos M. Opazo, Celeste H. Mawal, Alexandra Birrell, Joan So, Keziah Ting, Fiona Brown, Andrew H. Wei, Jason A. Powell, Kristin K. Brown, Ricky W. Johnstone, Lev M. Kats

## Abstract

The ubiquitous metabolite heme has diverse enzymatic and signalling functions in most mammalian cells. Cells can salvage heme from the extracellular environment or synthesise heme *de novo* from succinyl-CoA and glycine through a series of 8 enzymatic reactions catalysed by heme biosynthesis enzymes (HBEs) localised in the mitochondria and the cytosol^1,2^. Through integrated analyses of mouse models, human cell lines and primary patient samples, we identify *de novo* heme biosynthesis as a selective dependency in acute myeloid leukaemia (AML). The dependency is underpinned by a propensity of AML cells, and especially leukaemic stem cells (LSCs) to downregulate HBEs. The resultant low heme state upregulates self-renewal genes via the heme sensing transcription factor BACH1, but also places leukaemia cells on the threshold of heme starvation. Genetic or pharmacological inhibition of HBEs induces cuproptosis, a form of programmed cell death caused by copper accumulation and oligomerisation of lipoylated proteins^3^. Moreover, we identify pathways that are synthetic lethal with heme biosynthesis, including glycolysis, which can be leveraged for combination strategies. Altogether, our work uncovers a heme rheostat that controls gene expression and drug sensitivity in AML and implicates HBE inhibition as a novel cuproptosis trigger.

## Main

Dysregulated metabolism is a hallmark of cancer cells and contributes to disease maintenance and progression on multiple levels^4^. Many cancer driver genes including RAS, MYC and PTEN reprogram metabolic networks to produce energy and biomass required for unrestrained growth^5^. Metabolites such as α-ketoglutarate and S-adenosylmethionine are also required for epigenetic factors to control chromatin state and are thus intricately linked with aberrant self-renewal and differentiation in malignant cells^6^. Moreover, metabolic activity controls programmed cell death pathways, dictating whether tumour cells respond to stress through adaptation and survival or self-destruction^7^. Consequently, metabolic targeting can have potent anti-cancer effects, including in AML where metabolic inhibitors demonstrate clinical efficacy even in the relapsed/refractory setting^8–10^. Identification of additional tumour specific metabolic dependencies represents fertile ground for new drug development in areas of unmet need^11^.

Heme is an essential metabolite with broad biological activity. Beyond its function as an oxygen carrier in erythroid cells, heme is required for and regulates numerous molecular processes in non-erythroid cells ranging from mitochondrial energy generation to iron homeostasis, antioxidant defence, kinase signalling and transcription^1,2^. Heme levels have been shown to modulate cell fate decisions including differentiation^12,13^ and apoptotic cell death^14^. However, despite the potential for altered heme biosynthesis to impact on critical features of cancer cells, HBEs have not been extensively investigated as drug targets in cancer. Prompted by observations that HBE expression and heme levels are reduced in AML (our findings and Ref ^14^), combined with analysis of the Cancer Dependency Map identifying HBEs as highly selective AML dependencies (see below), led us to investigate the viability of HBE targeting as an anti-leukaemia strategy.

### Heme biosynthesis is downregulated in AML

To identify pathways that are required to sustain the transformed state of AML cells, we developed a novel multigenic mouse model (referred to as DIN) driven by a doxycycline inducible DNMT3A^R882H^ allele (one of the most common AML driver mutations)^15,16^. Genetic de-induction of DNMT3A^R882H^ in established leukaemic cells greatly increased survival of tumour bearing mice, indicating that mutant DNMT3A and its downstream effectors are required for disease maintenance *in vivo* (Extended Data Figs 1A-F). We sorted DIN progenitors from tumour-bearing mice following doxycycline treatment and scrutinised gene expression by RNA sequencing (RNAseq) (Fig 1A-B, Extended Data Figs 1G-I, Supplementary Table 1). Intriguingly, we found that one of the most prominent transcriptional responses to acute DNMT3A^R882H^ depletion was the upregulation of HBEs (seven out of eight enzymes significantly increased (Fig 1A-B), suggesting that AML oncogenes may downregulate HBEs. Indeed, a prior study that analysed patient microarray data reported that HBEs are broadly downregulated in AML and are among the most downregulated genes during disease progression^14^. To verify this observation in an independent patient cohort we analysed RNAseq data from BEAT AML^17^ using the rank based singscore single-sample scoring method^18,19^ to quantify HBE expression (Methods). Grouping patients by common recurrent mutations, we found that HBEs are generally expressed at lower levels in leukaemic cells compared with normal bone marrow cells, but we also noted significant heterogeneity among the different genetic groups (Fig 1C). Similar patterns (e.g. relatively high HBE expression in *TP53* and *CEBPA* mutant cases) were also observed in an additional Cancer Genome Atlas cohort although it lacks internal normal controls (Fig 1C)^15^. We also analysed a dataset that compared RNA and protein expression between haematopoietic stem/progenitor cells (HSPCs), LSCs and more mature AML blasts^20^, finding downregulation of heme metabolism in LSCs at the pathway and individual gene level (Fig 1D). Thus, HBE expression is downregulated in a considerable subset of AML and especially in AML LSCs.

**Figure 1.**
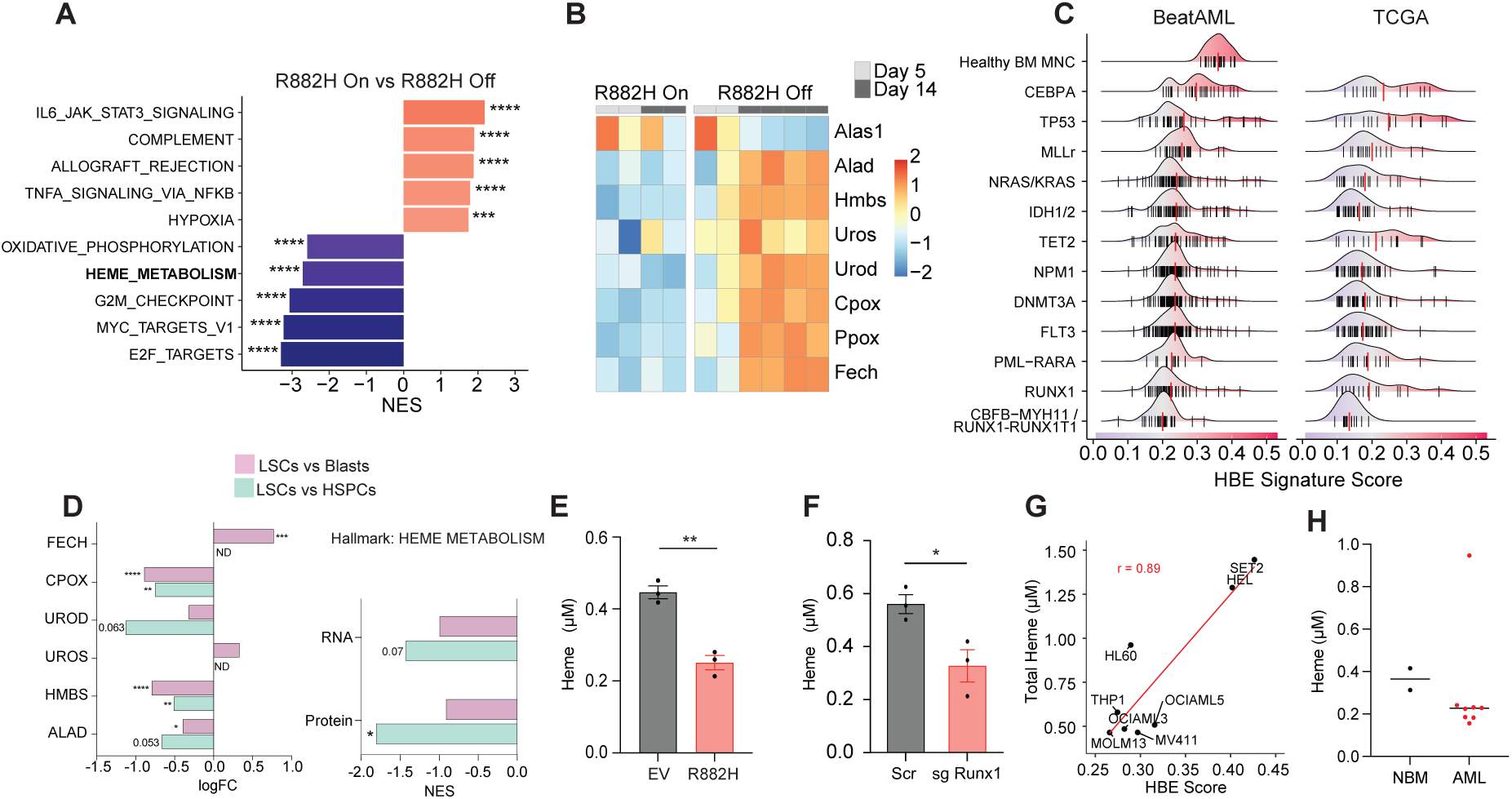
Heme biosynthesis is downregulated in AML. **A-B.** RNAseq analysis of cKit+CD11b-DIN progenitors from tumour bearing mice untreated (R882H on) or treated with doxycycline (R882H off) for 14 days (n = 2-4 mice/group). **A.** GSEA showing enrichment of selected signatures from the MSigDB database. * FDR < 0.05, ** FDR < 0.01, *** FDR < 0.001, **** FDR < 0.0001. **B.** Heatmap showing HBE expression. **C.** HBE scores calculated using the singscore method^18,19^ for AML patients and normal controls in the BEAT AML and TCGA cohorts. Each vertical line represents an individual patient. Patients are segregated based on the presence of the indicated recurrent genetic lesions. **D.** Transcriptomic and proteomic analysis of HBE expression in FLT3/NPM1 wild-type AMLs. LogFC, NES and significance (FDR) was reported by Raffel et al^20^. * FDR < 0.05, ** FDR < 0.01, *** FDR < 0.001, **** FDR < 0.0001, ND = No data. **E-F.** Total cellular heme quantified using the heme extraction assay in E HoxB8 EV and R882H or F HoxB8 Scrambled (Scr) and Runx1 KO cells. n = 3 biological replicates. Error bars represent mean ± SEM. Significance was assessed with an unpaired t-test ** p <0.01. **G.** Correlation plot between HBE score and total cellular heme in AML cell lines. (r represents Pearson correlation coefficient). **H.** Total heme levels in control bone marrow (NBM; n = 2) and bone marrow cells from AML patients (n = 9).

We next embarked on a set of experiments to examine the impact of co-ordinated reduced HBE expression on intracellular heme levels which had not been done previously. Expression of DNMT3A^R882H^ in Hoxb8 immortalised murine HSPCs (a model system extensively used to study the molecular and cellular effects of individual leukaemic drivers in an HSPC context^21–23^) induced gene expression changes similar to those observed *in vivo* in the DIN model and downregulated HBEs (Extended Data Figs 2A-C). Consistent with the known function of DNMT3A in promoting HSPC differentiation, DNMT3A^R882H^ Hoxb8 cells exhibited impaired myeloid differentiation in response to GM-CSF stimulation (Extended Data Fig 2D)^24^. Quantification of total cellular heme revealed a ∼50% reduction in DNMT3A^R882H^ Hoxb8 cells compared with controls (Fig 1E). Moreover, DNMT3A^R882H^ Hoxb8 cells demonstrated reduced capacity to convert 5-ALA (the natural substrate of the second enzyme in the pathway, ALAD) to protoporphyrin (PpIX; the penultimate metabolite) and delayed PpIX clearance in a 5-ALA pulse-chase assay, suggesting reduced heme synthesis pathway flux (Extended Data Figs 2E-G). To test the impact of a different AML mutation associated with reduced HBE expression we used CRISPR/Cas9 to delete the tumour suppressor RUNX1 which also reduced heme levels and pathway flux (Fig 1F, Extended Data Fig 2H).

We then analysed heme metabolism in a panel of human AML cell lines spanning a variety of disease subsets, genetic alterations, and HBE levels (Extended Data Figs 2I-K). We found a strong positive association between HBE expression and levels of total cellular heme (Fig 1G). An orthogonal method using a genetically encoded fluorescent heme biosensor^25^ similarly revealed that cell lines with high HBE expression possess high levels of labile heme (Extended Data Fig 2K). Finally, we compared total cellular heme levels in primary AML samples and bone marrow controls (collected from Burkitt lymphoma patients with no bone marrow involvement). 8 of the 9 AML samples analysed had reduced heme, although concordant with gene expression data there was significant variation (Fig 1H). Altogether, our data and that of earlier studies^14^ support the conclusion that reduced HBE expression and low heme levels are commonly shared features of AML tumours.

### Heme regulates self-renewal and metabolic gene expression programs in leukaemic cells

Heme has been implicated as a potent regulator of transcription^26–29^ leading us to investigate how altered heme levels in AML influence gene expression patterns. First, we focussed on our syngeneic Hoxb8 model and specifically on the heme sensing transcription factor BACH1 which was previously found to be required for LSC activity *in vivo* in a pooled RNA interference screen^30^. Heme binds to BACH1 via cysteine–proline motifs, promoting its displacement from chromatin and degradation via the ubiquitin proteasome system^26,31^.

Recruitment of Bach1 at chromatin (and total Bach1 proteins but not mRNA) was increased in DNMT3A^R882H^ Hoxb8 cells compared with control cells and was rescued by supplementation with hemin (a membrane permeable heme analogue)(Fig 2A, Extended Data Figs 3A-E, Supplementary Tables 2-3). To characterise gene expression programs that are regulated by the heme/BACH1 axis, we selected genes that were associated with heme sensitive Bach1 peaks and differentially expressed between DNMT3A^R882H^ and control Hoxb8 cells (Extended Data Fig 3F). Consistent with data from other cell types^32,33^ Bach1 binding was associated with both up and downregulated genes in DNMT3A^R882H^ Hoxb8 cells (Fig 2A). Among Bach1 bound DEGs we identified numerous genes implicated in cell fate decisions. Notably, key self-renewal genes (*Kit*, *Mef2c*, *Fos*, *Slpi* and *Zeb2*) were upregulated and conversely genes associated with differentiation (*Cebpa*, *Rxra*, *Csf1*, *Cd63* and *Gca*) were downregulated. Bach1 bound DEGs were also enriched for metabolic factors with metabolomic analyses and drug killing assays demonstrating concordant downstream phenotypes e.g. downregulation of *Slc7a11*, reduced L-cysteine and increased sensitivity to ferroptosis inducing agents RSL3 and APR-246^34,35^ in DNMT3A^R882H^ Hoxb8 cells (Extended Data Figs 3G-J, Supplementary Table 4). Hemin treatment reversed the gene expression changes of selected Bach1 target genes (Extended Data Fig 3K). Together these data suggest that a low heme/high BACH1 state promotes stemness and implicate HBE as a powerful modulator of therapeutic response across multiple pathways beyond BCL2 inhibition^14^.

**Figure 2.**
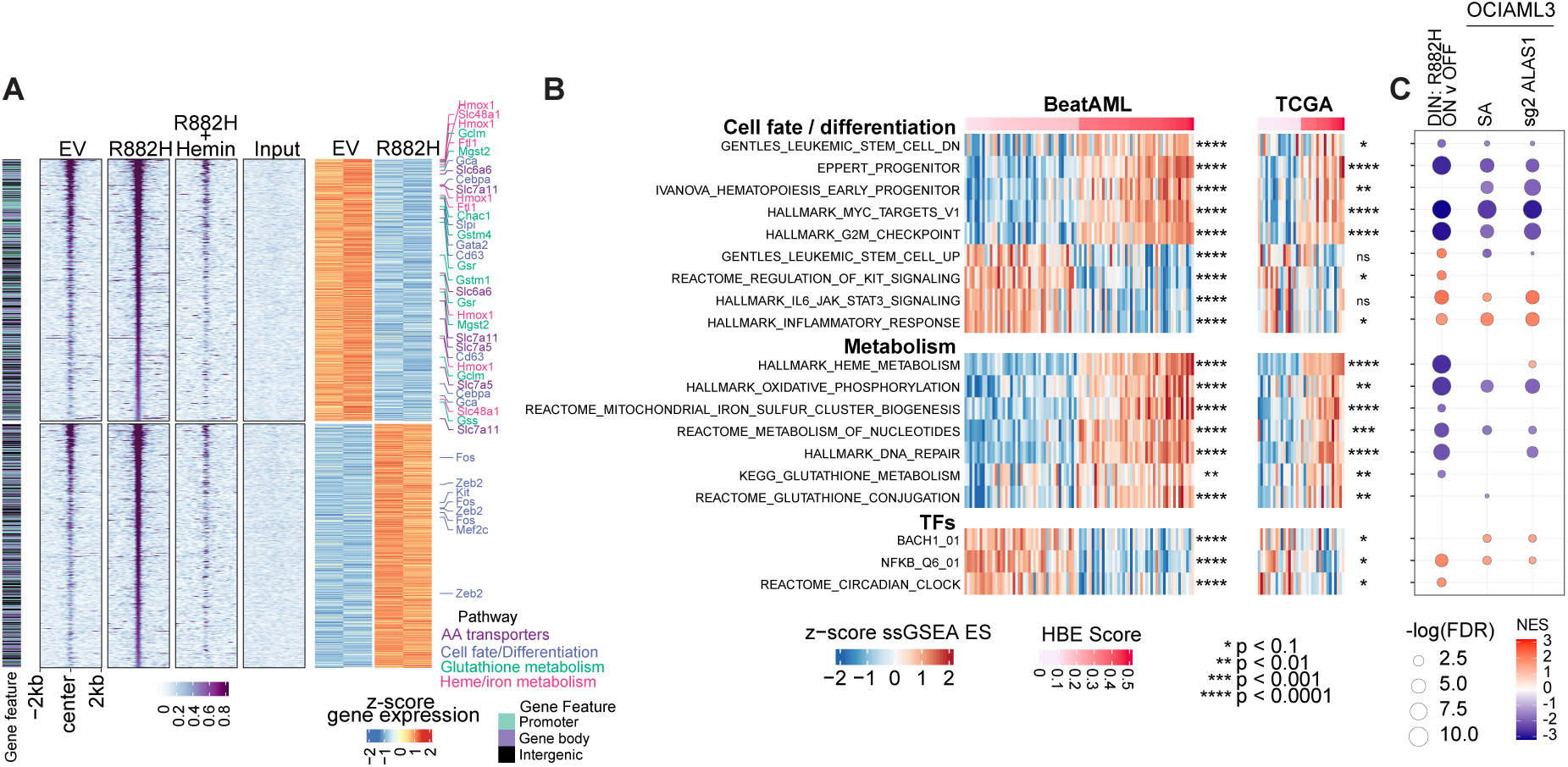
Heme regulates self-renewal and metabolic gene expression programs in leukaemic cells. **A.** Bach1 ChIPseq and RNAseq in Hoxb8 cells transduced with empty vector (EV) or DNMT3^AR882H^ treated with vehicle or hemin. Heatmap of Bach1 binding and gene expression at heme-responsive Bach1 bound peaks and their associated DEGs (overlap from Extended Data Fig. 3F). **B.** ssGSEA of the top and bottom 10% HBE high and low AMLs, resepctively, from the BEAT AML and TCGA cohorts showing enrichment of selected signatures. Wilcoxon test was used to assess the significance between the enrichment scores of HBE high and low AMLs. **C.** GSEA showing enrichment of signatures from B in the indicated conditions. Circles plotted when p < 0.15.

To further bolster our findings in Hoxb8 cells, we next analysed AML patient gene expression data. We ranked patients in the BEAT AML and TCGA cohorts based on their HBE score and then applied single-sample GSEA (ssGSEA)^36^ to compare signature enrichment from the Hallmarks, C2 and C3 collections of the MSigDB database between patients in the top (HBE high) and bottom (HBE low) 10% of HBE scores, respectively (Fig 2B). Consistent with observations in the Hoxb8 context, BACH1 and stemness signatures were upregulated and glutathione metabolism signatures were downregulated in HBE low patients. Interestingly, inflammation, circadian clock and NF-κB signatures were also preferentially enriched in HBE low patients whereas HBE high patients were enriched for signatures associated with high mitochondrial activity, myeloid differentiation and proliferation. The comparison between HBE high and low patients was further supported by an alternative methodology that correlated ssGSEA enrichment scores with HBE scores for all patients in the BEAT and TCGA cohorts (Extended Data Fig 3L). Heme is a known negative regulator of the circadian clock and represses the transcription factor CLOCK which is essential for LSC repopulating capacity^27,30,37^. Likewise, NF-κB activation has been described as a common feature of LSCs^38,39^. Correlation between low HBE expression and transcriptional signatures of LSCs and/or transcription factors known to be active in LSCs provides further evidence that LSCs are characterised by a low heme state.

For orthogonal validation, we tested the impact of directly modulating heme biosynthesis using genetic or pharmacological means in an AML cell line. We used the OCI-AML3 line which carries a DNMT3A mutation and is characterised by intermediate HBE expression and heme levels (Fig 1G, Extended Data Figs 2I, K). Deletion of ALAS1 or treatment with succinylacetone (SA; a tool inhibitor of ALAD widely used in pre-clinical studies^14,40^) reduced heme levels and recapitulated many of the transcriptional signatures evident in HBE low AML patients (Fig 2C, Extended Data Figs 3M-N, and Supplementary Table 5). The same signatures were also evident in the DIN model upon DNMT3A^R882H^ depletion (Fig 2C). Collectively, our data suggest that reduced heme levels drive altered metabolic and transcriptional programs that promote self-renewal of leukaemic cells but also give rise to therapeutic vulnerabilities.

### De novo heme biosynthesis is a selective leukaemia dependency

Reduced availability of, or increased demand for, essential metabolites in certain cancers are prone to generate enhanced reliance on specific pathways and manifest as targetable vulnerabilities^41–43^. Analysis of DepMap revealed heme biosynthesis as one of the most selectively dependent pathways in AML (Extended Data Figs 4A-B, Methods). All eight enzymes in the pathway have a mean negative dependency score in AML cell lines (indicative of an antiproliferative effect of the gene knockout), with scores for six out of eight enzymes being significantly lower in AML versus non-AML cell lines (Fig 3A). By comparison, genes implicated in heme salvage such as FLVCR2 and SLCO2B1 are not required for proliferation of AML cells (Extended Data Fig 4C). Next, we correlated the dependence on individual HBEs with HBE score across AML cells lines (Methods), finding that cell lines with low HBE score are especially dependent on HBEs (Fig 3B, Extended Data Fig 4D). Notably, cells with low HBE scores also demonstrate an increased dependence on BCL2 (and MCL1), consistent with the previously reported relationship between heme biosynthesis and intrinsic apoptosis pathways (Fig 3B)^14^.

**Figure 3:**
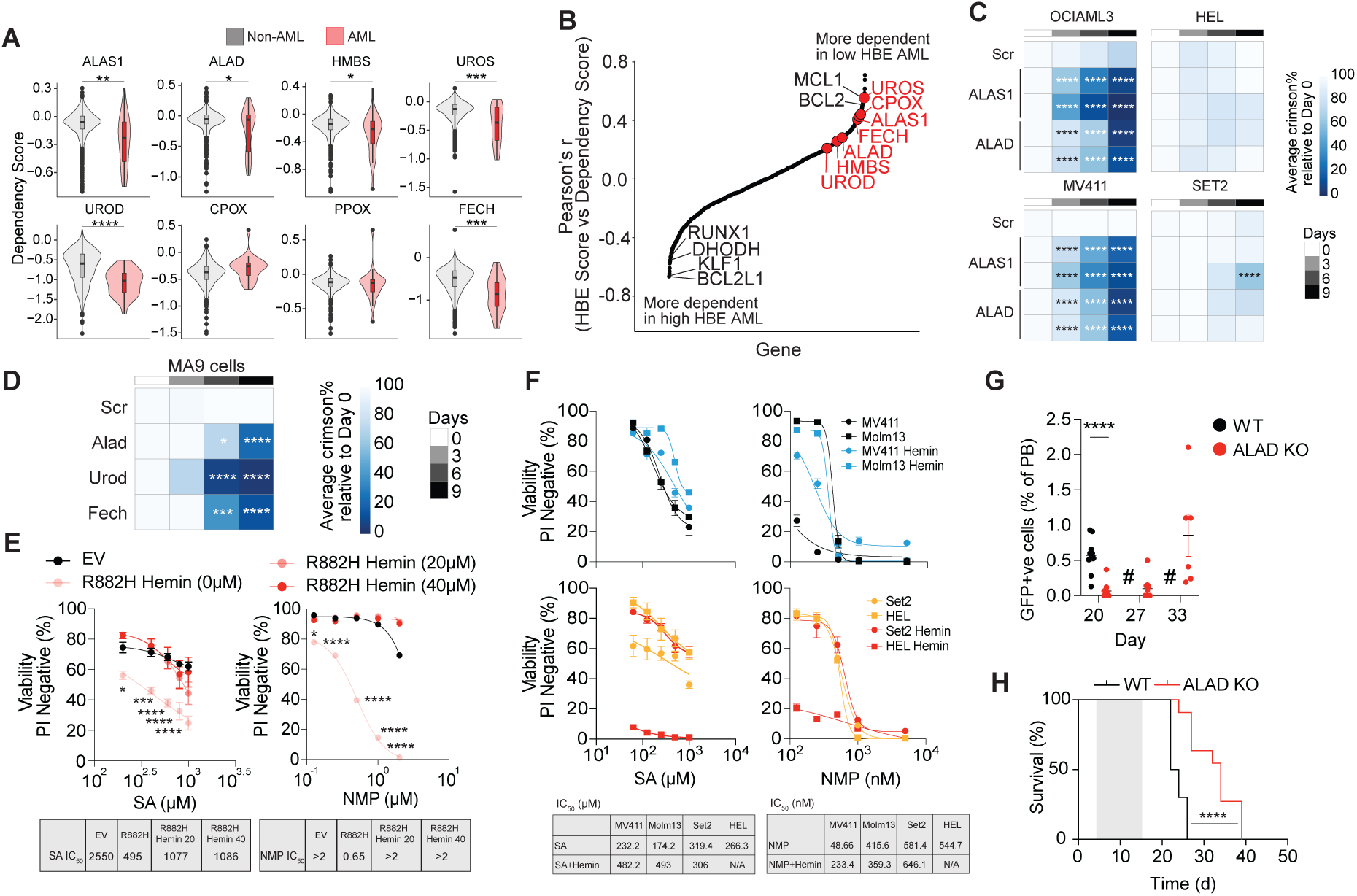
De novo heme biosynthesis is a selective leukaemia dependency. **A.** DepMap gene dependency scores for genes encoding HBEs comparing AML and non-AML cell lines. Significance was assessed with a one-sided t-test. * p < 0.05, ** p < 0.01, *** p < 0.001, **** p < 0.0001. **B.** Pearson’s correlation coefficient between HBE score vs dependency score in AML cell lines. **C.** Proliferative competition assay in AML cells transduced with sgRNAs targeting ALAS1, ALAD, UROD, FECH or control (Scr). Each column represents an independent sgRNA. n = 3 biological replicates. Significance was assessed by two-way ANOVA * p < 0.05, *** p < 0.001, **** p < 0.0001. **D.** Proliferative competition assay in primary murine MLL-AF9/Nras^G12D^ cells. n = 3 biological replicates. Significance was assessed by two-way ANOVA * p < 0.05, *** p < 0.001, **** p < 0.0001. **E-F.** Viability of **E.** Hoxb8 cells transduced with empty vector (EV) or DNMT3A^R882H^ **F.** or AML cell lines treated with SA or NMP in the presence of hemin. n = 3 biological replicates. Significance was assessed with a multiple comparison adjusted t-test; * p < 0.05, ** p < 0.01, *** p < 0.001, **** p < 0.0001. **G-H.** NSG mice engrafted with MV4-11 cells transduced with a doxycycline inducible ALAD sgRNA were treated with or without doxycycline. n = 10-11 mice/group. Shaded area denotes administration of doxycycline chow and water. **G.** Proportion of GFP+ AML cells in the peripheral blood at the indicated times quantified by flow cytometry. Error bars represent mean ± SEM. Significance was assessed with a two-sided t-test **** p < 0.0001. **H.** Kaplan-Meier survival curve. Significance was assessed by Mantel-Cox test **** p < 0.0001.

We performed proliferative competition assays targeting individual HBEs in two genetically distinct low/intermediate heme AML cell lines, OCI-AML3 and MV4-11. In line with the pooled CRISPR screening data, loss of HBEs resulted in significant proliferation defects associated with reduction in total cellular heme levels suggesting that heme import cannot compensate for the loss of *de novo* synthesis (Fig 3C, Extended Data Figs 4E-F). HBE deletion in primary murine AML cells derived from the aggressive MLL-AF9/Nras^G12D^ model exhibited similar effects (Fig 3D). By comparison, HBE targeting in high heme AML cell lines HEL and SET2 resulted in more muted phenotypes, notwithstanding comparable proliferation rates (Fig 3C).

To complement the genetic data we tested the anti-proliferative properties of two pharmacological inhibitors of HBE – SA and *N*-methylprotoporphyrin (NMP; an inhibitor of FECH^44^). We confirmed on target activity of SA and NMP, with SA blocking PpIX accumulation and NMP exacerbating PpIX accumulation upon 5-ALA supplementation as expected (Extended Data Fig 4G). Exposing Hoxb8 cells to SA or NMP induced cell death in a concentration-dependent manner with DNMT3A^R882H^ cells significantly more sensitive than isogeneic controls (Fig 3E). Notably, treatment with exogenous hemin reversed the increased sensitivity of DNMT3A^R882H^ cells, suggesting that SA and NMP mediated killing was attributable to on target activity of HBE inhibition (Fig 3E). In AML cell lines SA and NMP sensitivity similarly correlated with heme levels, with heme low cells (MV4-11 and MOLM13) more sensitive compared with heme high cells (HEL and SET2)(Fig 3F). Hemin supplementation also reduced but did not completely rescue the effects of HBE inhibition in AML cell lines with the exception of HEL cells where hemin induced anti-proliferative and pro-differentiation effects as previously described^45^. Altogether, these data are in agreeance with the hypothesis that reduced heme synthesis sensitises AML cells to HBE inhibition.

Genetic dependencies and especially metabolic dependencies may differ significantly *in vitro* and *in vivo*, in part because of different nutrient environments. To investigate the impact of disrupting heme biosynthesis in established leukaemias *in vivo*, we engrafted MV4-11 cells (widely used as a lethal disseminated AML model) expressing a doxycycline inducible ALAD guide and a constitutive GFP reporter into mice and treated them with doxycycline. Knocking out ALAD enabled us to track functional disruption of heme synthesis at single cell resolution using the 5-ALA assay. Within five days of guide induction, the capacity of AML cells to process 5-ALA into PpIX was abrogated in ∼70% of the cells (Extended Data Fig 4H). ALAD deletion significantly delayed disease development as evident by reduced circulating leukaemic cells in the peripheral blood and significantly prolonged survival of tumour bearing mice (Figs 3G-H). Of note, AML cells isolated from the ALAD knockout group at ethical end point demonstrated restoration of heme biosynthesis by the 5-ALA assay most likely due to the outgrowth of cells that had escaped ALAD deletion (Extended Data Figs 4H-I). Thus, *de novo* heme synthesis is required for AML growth *in vitro* and *in vivo*.

### Identification of metabolic modifiers of heme starvation by pooled CRISPR screening

Nutrient deprivation can activate diverse cell death pathways in a context dependent manner^46–48^. We devised a pooled CRISPR/Cas9 screen to delineate mechanism of cell death triggered by HBE inhibition and to uncover metabolic programs that are synthetic lethal with heme biosynthesis (Extended Data Fig 5A). OCI-AML3 cells transduced with a metabolomics focussed sgRNA library were cultured in the presence of DMSO (control), SA, NMP or hemin and next-generation sequencing used to compare positive and negative enrichment of gene knockouts in the various conditions (Methods)^49,50^. The hemin arm was included as a comparator that we hypothesised would reveal factors limiting HBE activity in AML.

Correlation of genes positively or negatively selected upon SA, NMP or hemin treatment to the control condition revealed a high concordance between the two HBE inhibitors, pointing to an overlap in the mechanisms that underpin their anti-proliferative effects (Extended Data Fig 5B). Notwithstanding the similarities between SA and NMP, there were also some differences with positive selection of genes more prominent with SA treatment and negative selection more pronounced with NMP treatment, especially at the later time point. This was likely the result of the relative anti-proliferative effects of the SA and NMP doses used in the screen and could also reflect differences in feedback mechanisms and secondary metabolic effects between ALAD and FECH inhibition. We examined genes involved in heme metabolism. The heme catabolism enzyme HMOX1 had no impact on proliferation in DMSO but was highly negatively selected upon hemin treatment consistent with an essential role in preventing heme-mediated toxicity (Extended Data Fig 5C). Conversely, and in line with DepMap data, HBE genes were depleted to varying degrees in DMSO (Extended Data Fig 5C). For most genes, the depletion was exacerbated with HBE inhibitor treatment and rescued by hemin (Extended Data Fig 5C). An exception to this expected pattern was UROD, which was severely depleted even in the context of hemin. We used proliferative competition assays with individual sgRNAs and confirmed the UROD phenotype across multiple AML cell lines (Extended Data Fig 5D). These data suggest that the strong antiproliferative effects of UROD deletion are at least partially independent of heme starvation. More broadly they confirm that our screen can identify heme reactive metabolic pathways.

### Heme starvation induces cuproptosis

Among the top hits that provided strong protection against the cytotoxic effects of HBE inhibition were genes required for the synthesis of lipoic acid (MECR, LIPT1, LIPT2, LIAS, DLD)(Fig 4A and Extended Data Fig 6A). Lipoylation is an evolutionary conserved posttranslational modification that is deposited on and required for a restricted subset of mitochondrial multi-protein complexes^51^. Importantly, genes encoding components of these complexes including GCSH, PDHA1 and PDHB were similarly positively selected in the screen specifically in the presence of SA and NMP (Fig 4A and Extended Data Fig 6A).

**Figure 4:**
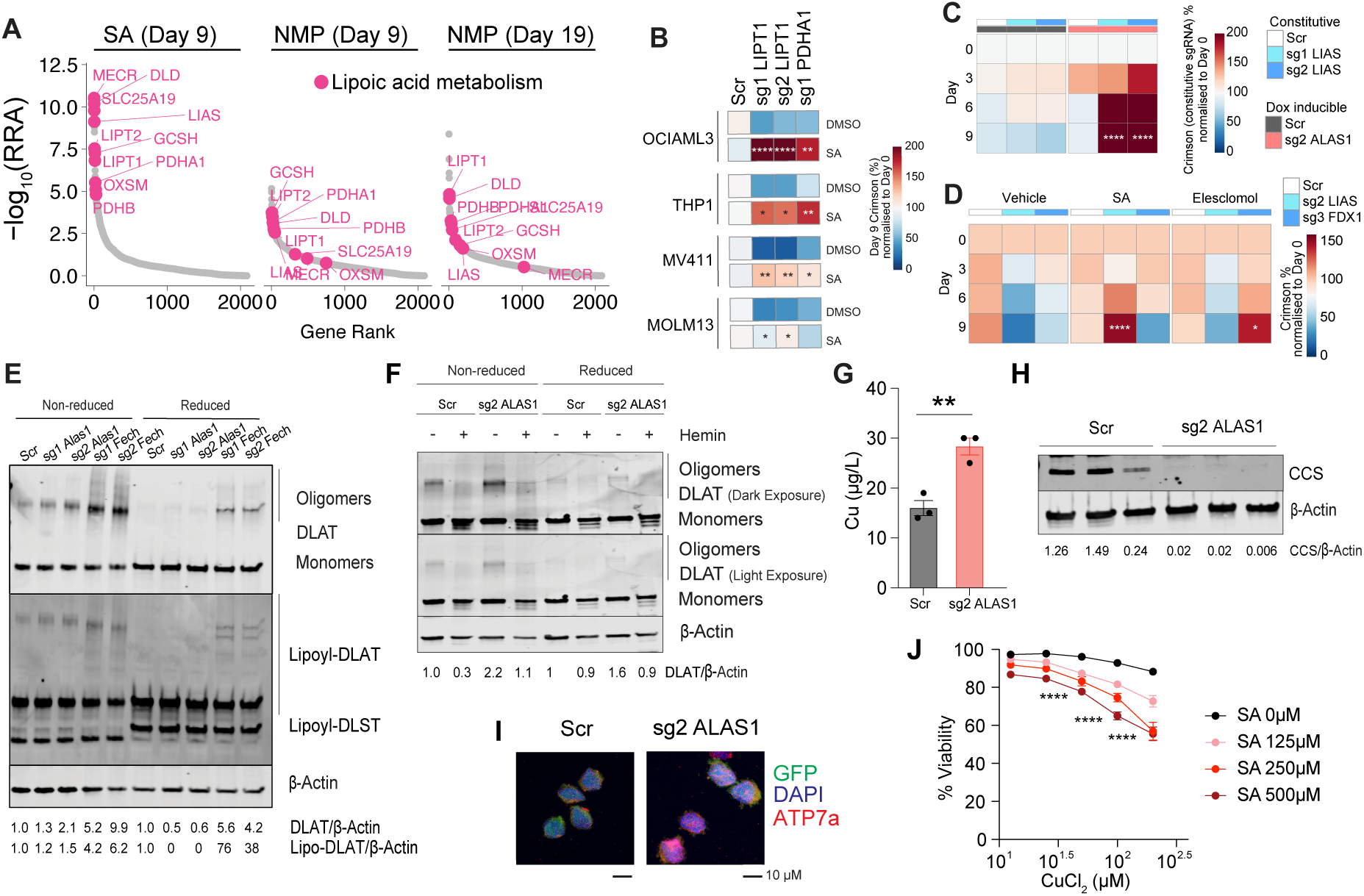
Heme starvation induces cuproptosis. **A.** MAGECK-RRA analysis identifying genes that decrease the sensitivity of AML cells to heme depletion by SA or NMP. **B.** Competition assays of AML cells transduced with sgRNAs targeting LIPT1 or PDHA1 and treated with SA. n = 2 biological replicates. Significance was assessed by two-way ANOVA * p <0.05, ** p <0.01, *** p <0.001, **** p <0.0001. **C.** Proliferative competition assays of OCI-AML3 cells transduced with doxycycline inducible scrambled or ALAS1 sgRNAs and constitutive LIAS sgRNA. n = 3 biological replicates. Significance was assessed by two-way ANOVA **** p <0.0001. **D.** Proliferative competition assays of OCI-AML3 cells transduced with scrambled, LIAS or FDX1 sgRNAs and treated with SA or elesclomol. n = 3 biological replicates. Significance was assessed by two-way ANOVA * p <0.05, **** p <0.0001. **E-F.** Western blot of OCI-AML3 cells transduced with doxycycline inducible sgRNAs and treated for 7 days with doxycycline to induce sgRNA expression and **F)** with or without hemin prior to analysis. **G.** Inductively coupled plasma mass spectrometry (ICP-MS) of copper levels in OCI-AML3 cells expressing ALAS1 or scrambled sgRNAs for 7 days. n = 3 biological replicates. Error bars represent mean ± SEM. Significance was assessed unpaired students T-test ** p < 0.01. **H.** Western blot analysis of copper chaperone CCS in OCI-AML3 cells expressing ALAS1 or scrambled sgRNAs for 7 days. **I.** Immunofluorescence of OCI-AML3 cells expressing doxycycline-inducible ALAS1 or scrambled sgRNAs stained with DAPI and anti-ATP7A antibody. **J.** Viability (PI negative) assays of OCI-AML3 cells treated with SA and copper chloride for 6 days. n = 3 biological replicates. Error bars represent mean ± SEM. Significance was assessed by two-way ANOVA * p <0.05, ** p <0.01, *** p <0.001, **** p <0.0001.

Lipoic acid synthesis was recently implicated as a key determinant of a novel form of programmed cell death caused by copper ionophores termed cuproptosis^3^. The striking overlap of cuproptosis resistance genes^3^ with our data prompted us to explore whether heme depletion causes cell death via similar mechanisms. Using competition assays we validated that knockout of LIPT1 or PDHA1 mediated resistance to HBE treatment in multiple AML cell lines (Fig 4B). To provide genetic evidence linking heme and lipoic acid synthesis pathways we generated doxycycline inducible ALAS1 knockout cells and performed competition assays with LIAS sgRNAs. As was the case with SA and NMP treatment, LIAS sgRNA expressing cells were positively enriched upon ALAS1 knockout (Fig 4C). Thus, disruption of lipoic acid synthesis provides protection against pharmacological or genetic inhibition of HBE.

The ferredoxin protein FDX1 is a master regulator of protein lipoylation^52^, but guides targeting FDX1 were absent from the targeted library used in our CRISPR screen. As expected, knockout of FDX1 induced resistance to the copper ionophore elesclomol^3^. In contrast however, FDX1 knockout sensitised AML cells to HBE inhibition with SA (Fig 4D). Aside from its function in lipoic acid synthesis, FDX1 is also required for conversion of heme *b* (the molecule synthesised by HBE) to heme *a* - a chemically modified form of heme that is covalently bound to components of the electron transport chain (ETC) such as cytochrome C1^1,53^. This requirement for FDX1 to produce heme *a* likely explains the additive effects of FDX1 knockout and SA treatment that we observe.

A hallmark of cuproptosis is the formation of reduction-sensitive oligomers of lipoylated proteins caused by chelation of the lipoic acid moiety by copper ions ^3^. ALAS1 and FECH knockout induced the formation of oligomers of the lipoylated protein DLAT (Fig 4E). The oligomers reacted not only with the anti-DLAT antibody, but also with an antibody directed against lipoic acid (Fig 4E). SA treatment, similarly, increased DLAT oligomerisation (Extended Data Fig 6B). Importantly, hemin supplementation reversed DLAT oligomerisation triggered by ALAS1 knockout and SA treatment, pointing to reduction in heme as the main driver of lipoylated protein aggregation (Fig 4F and Extended Data Fig 6C).

Although copper is an essential trace metal, intracellular copper concentration is actively maintained at miniscule levels in most cells to prevent copper toxicity. ALAS1 knockout and SA treatment increased intracellular copper levels by ∼75% as measured by inductively coupled plasma mass spectrometry (ICP-MS)(Fig 4G and Extended Data Fig 6D). As expected, we also observed a decrease in intracellular iron levels by ∼13% upon ALAS1 knockout (Extended Data Fig 6E). Concordant with the increase of intracellular copper we observed a decrease in the expression of the copper chaperone protein CCS in ALAS1 knockout cells, decreased CCS having previously been described as a marker of copper overload^54,55^ (Fig 4H). Similar decreases in CCS expression were observed using other genetic and pharmacological methods to reduce heme levels (Extended Data Fig 6F). Moreover, ALAS1 knockout also caused a marked intracellular elevation and redistribution away from the plasma membrane of the copper exporter ATP7A, an orthogonal indicator of increased intracellular copper^56^ (Fig 4I). Conversely, HBE inhibition did not alter expression of the copper importer SLC31A1 (Extended Data Fig 6G). Finally, supplementation of culture medium with copper significantly potentiated the effects of SA treatment and ALAS1 knockout, demonstrating the capacity of HBE inhibition to break copper homeostasis maintained by AML cells (Fig 4J and Extended Data Fig 6H). Altogether, our data demonstrate a direct and inverse relationship between heme and copper levels in AML cells and identify heme depletion as a potent trigger of cuproptosis.

### Synergistic effects of HBE and glycolysis inhibitors

We next analysed our screen for other dependencies that were influenced by HBE inhibition or heme supplementation. Top ranking hits that enhance the efficacy of NMP and SA were highly enriched for genes involved in glycolysis, including SLC2A1, PGP, GPI and GK (Fig 5A, Extended Data Fig 7A). Of note, these genes were dispensable under standard culture conditions and required only in the context of HBE inhibition. Individual gene knockouts of SLC2A1 and GPI robustly sensitised AML cells to SA and NMP treatment as expected (Fig 5B). These findings likely reflect the essentiality of heme as a co-factor for complexes III and IV of the ETC, and the synthetic lethal interaction between ETC and glycolysis components that has been demonstrated across different cellular contexts^57–60^. Other processes that were selected included sialic acid metabolism which was depleted upon SA and NMP treatment; and N-linked glycosylation (specifically mannosyltransferases) in the endoplasmic reticulum (ER) which was depleted upon hemin treatment (Extended Data Fig 7B-E). The latter potentially suggests that ER stress may be a barrier to increased heme production in AML cells.

**Figure 5.**
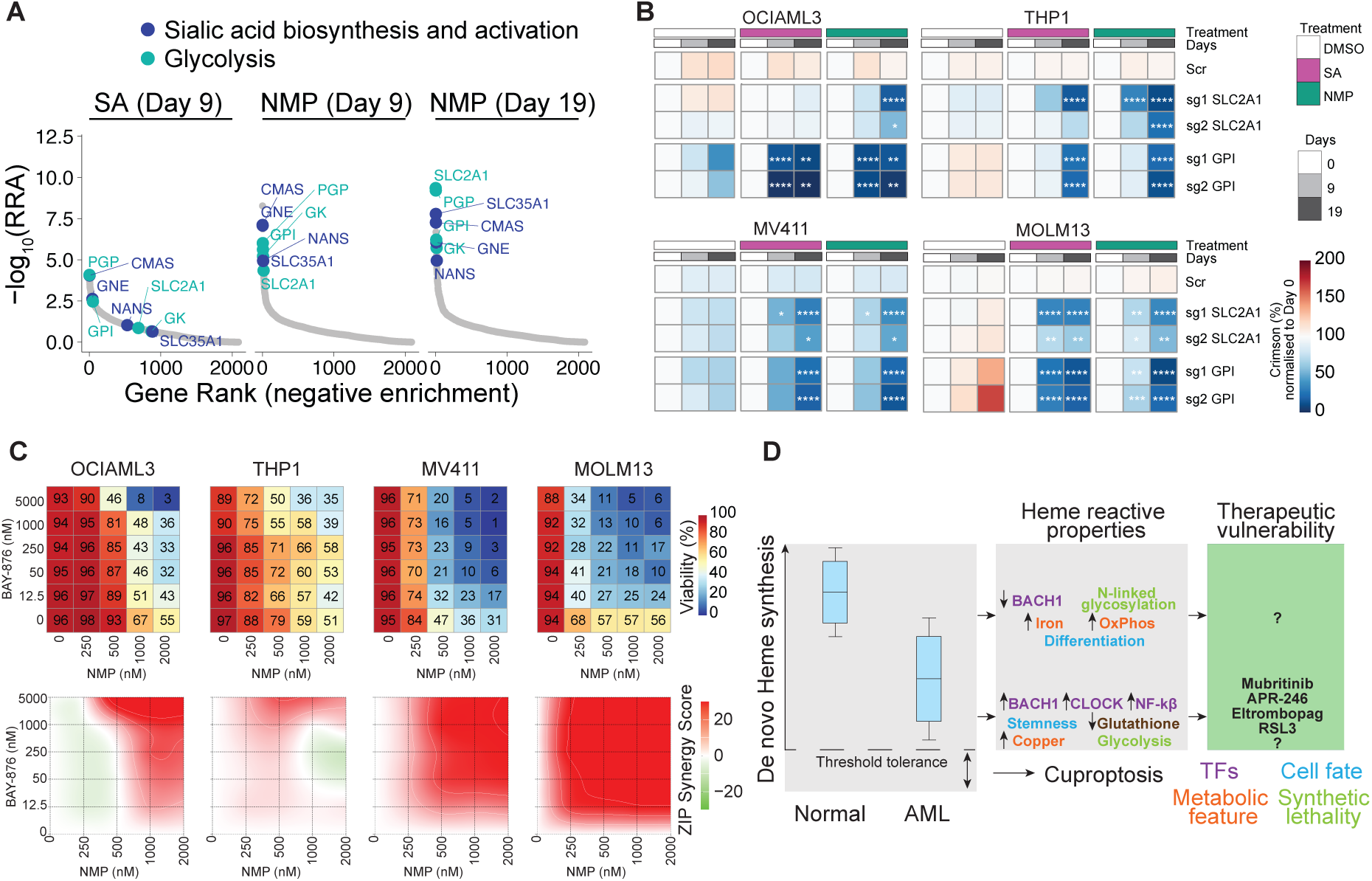
Synergistic effects of HBE and glycolysis inhibitors. **A.** MAGECK-RRA analysis identifying genes that increase the sensitivity of AML cells to heme depletion by SA or NMP. **B.** Proliferative competition assays of AML cells transduced with sgRNAs targeting SLC2A1 or GPI and treated with SA or NMP. n = 2 biological replicates. Significance was assessed by two-way ANOVA * p < 0.05, ** p < 0.01, *** p < 0.001, **** p <0.0001. **C.** Viability assays of AML cell lines treated with BAY-876 and NMP for 9 days. Drug synergy was determined using SynergyFinder (n = 3 biological replicates). **D.** AML cells have reduced de novo heme synthesis that contributes to transcriptional programs critical for cell growth. Heme dependent changes in transcriptional and metabolic targets can be exploited for therapeutic benefit. Reduced de novo heme synthesis and capacity of AML cells to buffer heme starvation above a critical threshold triggers cuproptosis upon HBE inhibition through increased copper import.

SLC2A1 (aka GLUT1) has emerged as a therapeutic target in various cancers, leading to the development of potent and selective inhibitors. We co-treated AML cell lines with NMP and the small molecule SLC2A1 inhibitor BAY-876^61^. Concordant with the genetic data we observed potent synergy between NMP and BAY-876 in all cell lines tested (Fig 5C). We also treated DNMT3A^R882H^ Hoxb8 cells with BAY-876, finding that they are significantly more sensitive to SLC2A1 inhibition than their isogenic counterparts (Extended Data Fig 7F). Interestingly, we noted the presence of a heme sensitive BACH1 peak associated with the *Slc2a1* locus in DNMT3A^R882H^ Hoxb8 cells (Supplementary Table 3). Although it did not pass our stringent statistical threshold for differential gene expression, there was a trend towards downregulation of Slc2a1 in DNMT3A^R882H^ Hoxb8 cells. Similar trends were observed in OCI-AML3 cells following treatment with SA or genetic ablation of ALAS1 (Supplementary Table 5). Moreover, we identified a strong positive correlation between HBE score and SLC2A1 expression in AML patient data (Extended Data Fig 7G). Together these observations suggest that the low heme/high BACH1 state of LSCs contributes to their inability to upregulate glycolysis in response to inhibition of mitochondrial respiration^62^. This feature of LSC metabolism differentiates LSCs from HSCs and is thought to be a crucial factor underpinning the efficacy of the venetoclax and azacitidine regimen that is becoming the standard-of-care for many AML patients^10^. Altogether, we have identified metabolic pathways that are functionally linked with heme production (Extended Data Fig 7H).

## Discussion

Heme is an abundant metallo-organic molecule required for diverse enzymatic and signalling functions. Almost all mammalian cells produce heme, but the regulation of heme metabolism, as well as the downstream consequences of altered heme availability remain poorly understood in most non-erythroid cell types^1,2^. In this study we demonstrate that *de novo* heme synthesis is a dependency in AML and correlates with reduced HBE expression. Notably, low HBE expression appears to be functionally important for AML due to a reciprocal relationship between heme and stemness, whereby AML LSCs downregulate HBE expression and the low heme state in turn promotes LSC activity. We identified BACH1 as a heme-sensing transcription factor that upregulates the expression of self-renewal genes in a low heme context. The connection between the low heme and LSC states is likely to be further reinforced by other heme sensing transcription factors such as CLOCK and potentially NF-κB^30,37–39^.

Acute heme depletion dysregulates copper homeostasis and triggers cuproptosis, a form of copper and lipoic acid dependent programmed cell death^3^. Given that cancer cells frequently inactivate programmed cell death pathways such as apoptosis, the discovery of molecularly independent cell death mechanisms lays the groundwork for development of novel therapies that can overcome drug resistance. Cuproptosis has emerged as one such pathway, although until now the only known cuproptosis trigger was copper overload by copper ionophores^3^. We implicate inhibition of heme biosynthesis as an independent cuproptosis trigger, further strengthening the notion that cuproptosis is a *bona fide* cell death pathway that may operate in a host of physiological and pathophysiological contexts. Of note, copper ionophores appear to be highly active against certain poor prognosis AML subtypes *in vitro*^63^.

Aside from severe depletion of heme leading to cell death, more moderate fluctuations in heme levels that alter cellular copper and/or iron levels also have the potential to impart profound phenotypes on haematopoietic cells and affect processes such as inflammation and HSC ageing^64,65^. Further research is needed to better understand the complex interconnected regulatory framework that maintains concentration of these molecules at the cellular and sub-cellular levels. Beyond metal ions, altered heme biosynthesis also impacts other metabolic networks with the potential for HBE expression and/or intracellular heme levels to serve as a biomarker for a range of clinically utilized and investigational agents such as BCL2 inhibitors, SLC2A1 inhibitors and ferroptosis activators^14,60,66,67^.

In conclusion, our study demonstrates that heme coordinates multiple fundamental aspects of AML biology and identifies HBEs as promising drug targets (Fig 5D). AML cells have an increased dependence on HBEs and reduced HBE expression sensitises cells to heme biosynthesis inhibitors. On the other hand, individuals with inherited hypomorphic mutations in HBE genes that severely reduce *de novo* heme production develop porphyria but survive to adulthood. Together, these observations point to a potential therapeutic window for drugs that block heme biosynthesis with the availability of HBE crystal structures and tool compounds providing a starting point for drug development.

## Acknowledgements

We thank the members of the molecular genomics, animal and flow cytometry core facilities at the Peter MacCallum Cancer Centre, Peter MacCallum Cancer Centre Models of Cancer Translational Research Centre and Metabolomics Australia at the University of Melbourne for technical support; Sharon La Fontaine, Jake Shortt and Mark Dawson for advice and critical review of the manuscript and Stuart Pitson for resource and financial support. We also thank our consumers advocates, Jane Weston and Sheree Dornan for their support of this work. This work was supported by grants from the Cancer Council of Victoria (APP1160018) and Impact Philanthropy. A.C.L. was supported by a fellowship from the Victorian Cancer Agency. E.G. was supported by a fellowship from Cure Cancer. M.D.J. was supported by the Australian Postgraduate Award.

## Author contributions

L.M.K, A.C.L. and E.G. designed the study; A.C.L, E.G., R.F., J.A., M.J.K, C.M.O., C.H.M, A.B., J.S. and L.M.K. performed the experiments and analysed the data; K.T., F.B., A.H.W., J.A.P, K.K.B. and R.W.J. provided technical expertise and essential reagents; A.C.L. E.G. and L.M.K wrote the manuscript; L.M.K. supervised the project.

## Methods

### Cell culture

AML cell lines were cultured in RPMI-1640 (Gibco) supplemented with 10% FBS, 1% penicillin/streptomycin at 37°C and 5% CO_2_. HEK293T cells were cultured in DMEM (Gibco) supplemented with 10% FBS and 1% penicillin/streptomycin. Hoxb8 lines were generated from Cas9-GFP knock-in mice (a generous gift from Marco Herold, Walter and Eliza Hall institute of Medical Research, Melbourne, Australia) as previously described^1^ and cultured in RPMI-1640 (Gibco) supplemented with 10% FBS, 100µM Glutamax, 50µM β-mercaptoethanol, 1% Penicillin/Streptomycin, 10µM β-estradiol and 35ng/ml hFLT3 ligand (POG complete media) at 37°C and 10% CO_2_. MA9-Cas9-Cherry cells^2^ were cultured *ex vivo* in low glucose DMEM (Invitrogen) supplemented with 20% FBS (Invitrogen), 0.1 mM l-Asparagine (Sigma-Aldrich), 1% penicillin/streptomycin (Invitrogen) at 37°C and 10% CO_2_. All cell lines were routinely tested for Mycoplasma contamination.

### Viral transduction

sgRNAs were cloned into lentiviral vectors lentiGuide-Crimson (Addgene plasmid #70683), lentiGuide-Puro (Addgene plasmid #52963) or FgH1tUTG-GFP (Addgene plasmid #70183). sgRNA sequences are provided in Table S7. Non-replicating lentiviruses were generated by transient co-transfection of the transfer plasmids into HEK293T cells together with the packaging plasmids pMDL (Addgene plasmid #12251), pRSV-REV (Addgene plasmid #12253), and pVSVG (Addgene plasmid #12259) using the calcium phosphate method. Transduction was performed by centrifuging target cells with virus containing media at 1000 x g in the presence of 4 µg/ml sequabrene (Sigma-Aldrich). All retroviral constructs were based on the MSCV backbone. The doxycycline (doxy)-inducible construct TRI-DNMT3A^R882H^ was generated by replacing the open reading frame encoding MLL-AF9 with an open reading frame encoding DNMT3A^R882H^ in an inducible MLL-AF9 vector^3^. MIT-NRAS^G12D^ was a kind gift from Johannes Zuber (Research Institute of Molecular Pathology, Vienna, Austria). To generate MIG-IDH2^R140Q^ the human IDH2^R140Q^ was cloned into MSCV-IRES-GFP (Addgene plasmid #20672). Retroviral generation and transduction of murine foetal liver cells was performed as previously described^4^.

### Flow cytometry and cell sorting

Cells were analysed on the BD LSR Fortessa X-20 or BD FACSymphony or sorted on the BD FACSAria Fusion 5 or BD FACSAria Fusion 3 (BD Biosciences) at the Peter MacCallum Cancer Centre flow cytometry core facility. Data was analysed using FlowJo (v 10.4). Cells were suspended in FACS buffer (2% FBS in PBS) and stained with fluorophore-conjugated antibodies where indicated. For peripheral blood, bone marrow or spleens from mice, red blood cell lysis was performed by incubating samples in ACK red cell lysis buffer (150mM NH_4_Cl, 10mM KHCO_3_, 0.1mM EDTA) for 2 min. Cells were then washed in FACS buffer and re-suspended in FACS buffer prior to analysis or staining.

### Animal experimental models

All animal experiments were performed at the Peter MacCallum Cancer Centre in an approved premises nominated on the Bureau of Animal Welfare Scientific Licence SPPL20183 (Agriculture Victoria, Australia) and were approved by the Peter MacCallum Cancer Centre Animal Experimentation Ethics Committee. C57BL/6, congenic C57BL/6.SJLPtprca (referred to as *Ptprca*) and immune deficient NOD.Cg-Prkdcscid Il2rgtm1Wjl/SzJ (referred to as NSG) mice were bred within the Peter MacCallum Cancer Centre animal facility or purchased from the Walter and Eliza Hall Institute of Medical Research (Melbourne, Australia). All experimental mice were housed at the Peter MacCallum Cancer Centre animal facility under specific pathogen-free conditions. Animals were group-housed in individually ventilated micro-isolator cages (6 mice per cage) on a 13hr light/11-hr dark cycle. Mice had continuous access to sterilised water and Barastoc irradiated mouse cubes (Ridley AgriProducts).

### Conditional deletion of ALAD in vivo

5x10^5^ MV4-11-Cas9-Cherry cells transduced with sg2 ALAD expressed from the FgH1tUTG-GFP lentiviral vector were transplanted into NSG mice via tail vein injection. Following an initial engraftment period mice were randomised to receive or not doxycycline chow (2500 mg/kg) and water (2 mg/mL). Animals were closely monitored for clinical signs of disease development (weight loss, lethargy, hunched posture) by technical staff who were blinded as to the conditions of the experiment. Peripheral blood was routinely collected by tail vein incision into EDTA coated Microvette capillary blood collection tubes (Sarstedt AG & Co) and leukaemia burden quantified by flow cytometry using GFP and hCD45 (PE-Cy7 mouse anti-human CD45 (BD Pharmingen #557748)) expression. Animals were euthanized at ethical endpoint based on clinical symptoms and overall survival rates were assessed using Kaplan-Meier analysis. Editing efficiency was quantified 5 days after doxycycline administration or at the ethical end point using the 5-ALA assay. For the latter, cryopreserved samples from all mice were thawed at the same time and cultured *ex vivo* overnight prior to the assay being performed.

### DIN mouse model

The DIN model was generated by transplanting foetal liver cells co-transduced with TRI-DNMT3A^R882H^, MIG-IDH2^R140Q^ and MIT-NRAS^G12D^ into sub-lethally irradiated (5.5Gy whole body irradiation delivered by an X-RAD 320 (Precision) *Ptprca* recipients as previously described^4^. For serial transplantation experiments, bone marrow or spleen cells from moribund animals were collected and cryopreserved in FBS supplemented with 10% (v/v) DMSO. Cryopreserved cells were thawed and washed in PBS and transplanted into sub-lethally irradiated *Ptprca* recipients. Disease monitoring and survival analysis were performed as above except using GFP and dsRed expression and staining with c-Kit (APC rat anti-mouse CD117 (BD Pharmingen #553356)) and CD11b (V450 rat anti CD11b (BD Biosciences #560455)). Deinduction of DNMT3A^R882H^ was achieved by administering doxycycline chow and water as above.

### Primary AML samples

Bone marrow and peripheral blood samples were collected from patients treated at the Peter MacCallum Cancer Centre and the Alfred Hospital who had newly diagnosed or morphologically relapsed AML. Samples were collected after informed consent and studies were conducted in accordance with the approved protocols (project 21/26) administered by Human Research Ethics Committee of the Walter and Eliza Hall Institute of Medical Research. Cryopreserved AML samples from diagnosis containing >65% blasts were thawed dropwise in IMDM containing 20% FBS (thaw media) and incubated at RT for 15min. Cells were pelleted by centrifugation at 1700rpm for 5min, washed in thaw media and centrifuged at 1700rpm for 5min. Cells were resuspended in IMDM with 10% FBS and counted for further analysis. Bone marrow aspirates from patients diagnosed with Burkitt’s lymphoma with no bone marrow involvement were used as a normal bone marrow comparison. Details of patient samples are provided in Table S8.

### Targeted pooled CRISPR screen

We designed a metabolism focussed sgRNA library comprised of 12666 sgRNAs targeting 2068 genes (6 sgRNAs/gene) directly involved in metabolism and 150 non-targeting negative control sgRNAs. sgRNA sequences were adapted from the human CRISPR metabolic gene knockout library (Addgene pooled library #110066)^5^ and included genes from the Metabolic Atlas (Human-GEM v1.3.0), Mammalian Metabolic Enzyme Database^6^ and members of the Solute Carrier (SLC) family. The library was cloned into the lentiGuide-Puro lentiviral vector (Addgene plasmid #52963) using Golden Gate cloning. sgRNA sequences are provided in Table S9. The screen was performed using standard protocols^7^ with modifications. sgRNA library virus was produced by transfection of HEK293T cells. OCI-AML3-Cas9-Cherry cells were transduced at an MOI < 0.3 at an average 1000-fold representation. Two days after transduction, cells were selected by puromycin (1µg/ml) treatment for 5 days. A T0 cell pellet was collected by centrifuging 1.2x10^7^ transduced OCI-AML3-Cas9-Cherry cells, snap frozen and stored at –80°C. Puromycin selected cells were split into 4 conditions: vehicle (1:1250 DMSO), succinylacetone (250µM), NMP (500nM) and hemin (20µM). Cells were split and drugs/vehicle were replenished every 3-4 days, maintaining a minimum of 1.2x10^7^ cells for each condition. Cell pellets from each condition were collected at day 9 stored at –80°C. For NMP and hemin, drug concentrations were increased to 750nM and 40µM, respectively, with additional pellets collected on day 19. Genomic DNA was extracted using the DNeasy Blood and Tissue Kit (Qiagen), libraries were prepared by nested PCR^7^, pooled and purified using AMPure XP beads (Beckman Coulter), and sequenced to a depth of ∼12 million reads with single-end 75 bp sequencing on a NextSeq500 (Illumina). Sequencing reads were demultiplexed using bcl2fastq (v2.17.1.14), and low-quality reads Q < 30 were removed. The reads were trimmed using cutadapt (v1.14)^8^ to extract the 20 bp targeting sequence and sgRNAs. The MAGeCK RRA algorithm (v0.5.8.1) (count and test) were used to quantify, normalise and identify genes that were enriched or depleted in the various conditions^9^. R packages ggplot2 (v 3.3.3) and pheatmap (v 1.0.12) were used for figure generation.

### Competitive proliferation assays

Cell lines expressing Cas9 were transduced with sgRNA constructs in the lentiGuide-Crimson (Addgene plasmid #70683) or FgH1tUTG-GFP (Addgene plasmid #70183) backbones with partial transduction efficiency (10-70%). Following FgH1tUTG-GFP transduction, cells were treated with doxycycline (1µg/ml) for 4 days to induce sgRNA expression prior to beginning the assay. Where indicated, OCI-AML3, MV4-11, MOLM13 and THP1 cells were treated with vehicle (1:1000 DMSO), SA (250μM, 50μM, 40μM or 250μM, respectively), NMP (500nM, 300nM, 200nM or 400nM, respectively) or hemin (OCI-AML3 only 20μM, 40μM) beginning on day 4 post transduction. Cells were treated with various compounds and passaged every 2-4 days. The percentage of sgRNA expressing cells (Crimson^+^ or GFP^+^) was measured by flow cytometry and normalized to the percentage at T0 (i.e. 3 days post transduction or 4 days post doxycycline treatment).

### Viability assays

2x10^4^ Hoxb8 cells or AML cell line cells were seeded in technical duplicate for each condition in a 96-well plate and treated with various compounds. Cells were passaged and drugs were refreshed every 2-3 days. Viable cells were quantified by flow cytometry using propidium iodide or DAPI exclusion. Cell numbers were calculated using CountBright Plus Absolute Counting Beads.

### qPCR

HoxB8 EV and DNMT3A^R882H^ cells were treated for 24hrs with DMSO or hemin (40µM). RNA was extracted using the Direct-zol RNA Miniprep kit (Zymo Research) according to the manufacturer’s instructions. qPCR was performed using SensiFast SYBR Hi-ROX kit (Bioline) on a CFX96 Touch Real-Time PCR System (Bio-Rad). Results were analysed using the ΔΔC(t) method and normalised to β-actin. qPCR primers sequences are provided in Table S7

### Western Blotting

Western blotting was performed according to standard laboratory protocols. Cells were lysed in RIPA lysis buffer (50 mM Tris–HCl pH 8, 150 mM NaCl, 1% (v/v) NP-40, 0.5% (v/v) sodium deoxycholate, 0.1% (w/v) SDS, 0.5 U benzonase nuclease, cOmplete mini protease inhibitor cocktail (Sigma-Aldrich) for 30 min at 4°C. Protein concentration was determined using Pierce BCA Protein Assay Kit (Thermo Fisher). Samples were then split into reduced and non-reduced conditions. For non-reduced samples 4x non-reducing sample buffer (8% (w/v) SDS, 40% (v/v) glycerol, 100mM Tris-HCl pH 6.8, 0.01% (w/v) Bromophenol Blue) was added. For reduced samples, 4x reducing sample buffer (8% (w/v) SDS, 40% (v/v) glycerol, 10% (v/v) β-mercaptoethanol, 100mM Tris-HCl pH 6.8, 0.01% (w/v) Bromophenol Blue) was added and samples were incubated at 95°C for 5min. Protein lysates were resolved on Mini-PROTEAN TGX Precast Protein Gels (Bio-Rad) and immunoblotted onto Immobilon-FL PVDF membrane 0.45 μm (Merck) or Nitrocellulose membrane 0.45 μm (Thermo Fisher).

Membranes were incubated with primary and secondary antibodies, and near-infrared Western blot detection was performed by the Odyssey CLx imaging system (LI-COR Biosciences). The following antibodies were used: DNMT3A (D23G1) rabbit mAb (1:1000), (Cell Signaling #3598), lipoic acid rabbit mAb (1:1000)(Abcam #ab58724), β-actin (1:5000) (Sigma-Aldrich #A2228), DLAT (4A4-B6-C10) mouse mAb (1:1000)(Cell Signaling, #12362), Bach1 rabbit pAb (1:500 - 1:1000)(Proteintech, 14018-1-AP), IRDye 680RD goat anti-Mouse IgG (H + L) (1:20000 - 1:40000)(LI-COR Biosciences #926-68070), IRDye 800CW goat anti-rabbit IgG (H + L) (1:20000 – 1:40000)(LI-COR Biosciences #926-32211).

### Immunofluorescence

OCI-AML3-Cas9 sg Scr and sg2 ALAS1 cells were treated with doxycycline (1 μg/ml) for 7 days to induce sgRNA expression. Microscope Cover Glass No. 1.5H, High Precision 10mm Diameter coverslips were coated in 0.1mg/ml polylysine 24hrs prior to plating cells. Coverslips were washed twice in sterile PBS and 50000 OCI-AML3-Cas9 sg Scr or sg2 ALAS1 cells were added to individual wells. Cells were allowed to settle at the bottom of the coverslip for 24hrs before fixing with 4% paraformaldehyde (PFA) for 15mins at room temperature. PFA was removed by aspiration, followed by rinsing with PBS and x3 5min washes with PBS. Coverslips were incubated with permeabilization buffer (0.1% Triton x-100 in PBS) for 15mins. Permeabilization buffer was removed by aspiration followed by rinsing with PBS and x3 5min washes with PBS. Coverslips were blocked with 1% (w/v) BSA in PBS overnight at 4°C. The following day, blocking buffer was aspirated followed by rinsing with PBS and x3 5min washes with PBS. Coverslips were incubated with mouse anti-human ATP7a (R1) antibody for 1hr at room temperature prior to aspiration followed by rinsing with PBS and x3 5min washes with PBS. Coverslips were incubated with anti-mouse-AF647 secondary antibody for 1hr at room temperature in the dark prior to aspiration followed by rinsing with PBS and x3 5min washes with PBS. Nuclei were stained with DAPI (1:10000 in PBS) for 5mins. Coverslips were washed x3 in PBS and dabbed dry to remove excess liquid and mounted with ProLong Diamond mounting media on a glass slide. Coverslip was dried for at least 24hr in the dark prior to imaging.

### Metabolomic analysis

Metabolomic analysis was performed at the Metabolomics Australia facility, University of Melbourne. A monophasic extraction protocol was used to extract the metabolites from Hoxb8 cells. To the cells, 250 µL of chilled 1:3:1 Chloroform/MeOH/Milli-Q water containing 0.5 nmol of ^13^C5_15_N valine and 0.5 nmol of ^13^C_6_ sorbitol was added. Each sample was vortexed and then incubated at 4°C for 10 min with continuous agitation (950 rpm) using an Eppendorf Thermomixer C. The samples were centrifuged at 4°C for 10 min at 12700 rpm using an Eppendorf centrifuge 5430 R. The supernatant was transferred into a fresh 1.5 mL Eppendorf tube and the cell debris was discarded. For GCMS analysis, a 50 µL aliquot of each sample was pooled to create the pooled biological quality control (PBQC). 50 µL of each study sample and the PBQC were transferred into HPLC inserts and evaporated at 30 °C to complete dryness, using a CHRIST RVC 2-33 CD plus speed vacuum. To limit the amount of moisture, present in the insert, 50 µL 100% methanol (LCMS grade) was added to each insert and evaporated using a speed vacuum.

Dried samples for targeted analysis were derivatised online using the Shimadzu AOC6000 autosampler robot. Derivatisation was achieved by the addition of 25 µL methoxyamine hydrochloride (30 mg/mL in pyridine, Merck) followed by shaking at 37°C for 2h. Samples were then derivatised with 25 µL of N,O-bis (trimethylsilyl)trifluoroacetamide with trimethylchlorosilane (BSTFA with 1% TMCS, Thermo Scientific) for 1h at 37°C. The sample was allowed to equilibrate at room temperature for 1 h before 1 µL was injected onto the GC column using a hot needle technique. Split (1:10) injections were performed for each sample. The GC-MS system used comprised of an AOC6000 autosampler, a 2030 Shimadzu gas chromatograph and a 50NX triple quadrupole mass spectrometer (Shimadzu, Japan) with an electron ionisation source(−70eV). The mass spectrometer was tuned according to the manufacturer’s recommendations using tris-(perfluorobutyl)-amine (CF43). GC-MS was performed on a 30m Agilent DB-5 column with 0.25mm internal diameter column and 1µm film thickness. The injection temperature (inlet) was set at 280°C, the MS transfer line at 280°C and the ion source adjusted to 200°C. Helium was used as the carrier gas at a flow rate of 1 mL/min and argon gas was used in the collision cell to generate the MRM product ion. The analysis of the derivatised samples was performed under the following oven temperature program; 100°C start temperature, hold for 4 minutes, followed by a 10°C/min oven temperature ramp to 320°C with a following final hold for 11 minutes. Samples were analysed in a randomised order, with a pooled biological quality control sample added after every 5 samples. The pooled biological quality control was used to monitor downstream sample stability and analytical reproducibility. Approximately 598 targets were collected using the Shimadzu Smart Metabolite Database, where each target comprised a quantifier MRM along with a qualifier MRM, which covers approximately 381 unique endogenous metabolites and multiple stable isotopically labelled internal standards. Resultant data was processed using Shimadzu LabSolutions Insight software, where peak integrations were visually validated and manually corrected where required. The dataset was normalised by median and log transformed.

### RNAseq and analysis of publicly available gene expression data

Cells were lysed in Trizol reagent (Thermo Fisher) and RNA was extracted using the Direct-zol RNA Miniprep kit (Zymo Research). RNAseq was performed at the Peter MacCallum Cancer Centre Molecular Genomics core facility. RNAseq libraries were generated using the QuantSeq 3′ mRNA-seq Library Prep Kit for Illumina (Lexogen) and sequenced on an Illumina NextSeq500 to produce 75-100 bp single-end reads with a depth of 6-10x10^6^ reads per sample. Sequencing reads were de-multiplexed using bcl2fastq (v2.17.1.14), low quality reads (Q < 30) removed and trimmed using cutadapt (v1.14)^8^. Reads were mapped to the mouse (mm10) or human (hg19) reference genomes using HISAT2 (v2.0.4)^10^ and counted using featureCounts (- O) from subread (v 2.0.0)^11^. Counts were filtered using filterByExpr from edgeR (v 3.34.0)^12^ with default parameters, quantile normalised using voom and differential gene expression performed using Limma (v 3.48.3)^13^. Gene set enrichment analysis was performed using fgsea (v.1.16.0)^14^ using the MSigDB (v2023.2) gene sets.

CCLE normalised counts (21Q4) were downloaded from DepMap portal (https://depmap.org/portal/). TCGA batch corrected and normalised RNAseq data was downloaded from the Pan Cancer Atlas^15^. BeatAML RPKM gene expression of AML patients was downloaded using AMLbeatR (v 0.1.0) and used for figure generation when normal samples were not needed. When comparing AML to healthy samples, BeatAML rawcounts for all patients and healthy samples was downloaded from vizome (http://www.vizome.org/additional_figures_BeatAML.html) and normalised as previously described^16^. Briefly, genes were removed if they had duplicate gene symbols or if the counts were <10 for 90% or more of the samples. Samples were removed if they were excluded from the RNAseq analysis by BeatAML (rnaSeqAnalysis = n in the Clinical Summary table). GC-content-corrected log_2_ RPKM were generated using conditional quantile normalization cqn (v 1.36.0)^17^.

Singscore (v 1.10.0)^18,19^ was used to calculate signature scores of HBEs (ALAD, HMBS, UROS, UROD, CPOX, PPOX, FECH) using the normalised gene expression counts. This signature score is referred to as HBE score in the text.

AML selective dependency scores for each gene were calculated by subtracting the average DepMap dependency score of non-AML cell lines from AML cell lines.

To correlate HBE score with Dependency scores from AML cell lines in the CCLE database, HBE score was first determined using the singscore method on CCLE tpm data after filtering for genes that were expressed at tpm > 1 in at least one AML cell lines. HBE score was then correlated with the dependency score of every gene. Genes were removed if they were not dependent in at least 4 AML cell lines (dependency score > -0.5) or it they were highly dependent in every AML cell line (dependency score < -0.8).

Figures were generated in R (v 4.0.2) using ggplot2 (v 3.3.3), ggridges (v 0.5.3), pheatmap (v 1.0.12) and ComplexHeatmap (v 2.6.2)^20^. Limma (v 3.46.0) was used to generate barcodeplots and perform rotating gene set testing.

### ChIPseq

1.5x10^7^ HoxB8 EV and DNMT3A^R882H^ cells were seeded at 5x10^5^ per ml in POG complete media. DNMT3A^R882H^ cells were treated with DMSO (0.16% v/v) or hemin (40μM) for 24hrs. Samples were fixed with formaldehyde solution (1% w/v) for 15mins with gentle rocking and quenched by adding 2.5M glycine for 5mins. 2x10^7^ cells were washed twice in ice cold PBS and resuspended in 1ml of ChIP Lysis buffer (1% SDS (w/v), 10mM EDTA, 50mM Tris-HCl pH 8.0, cOmplete mini EDTA-free protease inhibitor cocktail (Sigma-Aldrich)). Each sample was sonicated using a Covaris S220 Focused Ultrasonicator for 40 cycles (30s on (5% duty cycle; intensity - 5200 cycles per burst), 20s off). 9ml of modified RIPA buffer (1% (v/v) TritonX100, 0.1% (v/v) deoxycholate, 90mM NaCl, 10mM Tris-HCl pH8, cOmplete mini EDTA-free protease inhibitor cocktail (Sigma-Aldrich)) was added to each sample. 5% from each sample was collected and pooled for an input sample. 2.5µg of Bach1 antibody (R&D systems #AF5777) was added to each sample with gentle rocking overnight at 4°C. Protein G Dynabeads (Theromo Fisher) were washed twice in modified RIPA buffer and added to each sample for 2hrs with gentle mixing at 4°C. Dynabeads were washed 3 times in wash buffer (0.1% (w/v) SDS, 1% (v/v) TritonX100 v/v, 2mM EDTA, 150mM NaCl, 20mM Tris-HCl pH 8) for 5 min. Reverse crosslinking buffer: (1% (w/v) SDS, 100mM NaHCO_3_, 200mM NaCl, 300µg/ml Proteinase K) was added to each sample and incubated at 65°C for overnight. DNA was extracted with ChIP DNA Clean & Concentrator kit (Zymo Research). Sequencing libraries were prepared using NEBNext Ultra II DNA Library Prep Kit for Illumina kit (New England Biolabs) according to the manufacturer’s protocol. Cleanup of adaptor ligated DNA was performed using AMPure XP beads. Purified DNA was amplified using PCR enrichment with 12 and 16 cycles for input and ChIP samples respectively. Libraries were sequenced on an Illumina Nextseq500 with 75bp single end reads to a depth of 21-35 million reads.

Raw sequencing reads were demultiplexed using bcl2fastq (v2.17.1.14) and adaptor sequences were removed using Trim Galore (-q 30) (v0.4.4) and reads aligned to the mouse genome (mm10) using Bowtie2 (v2.3.3) using default parameters^21,22^. Samtools (v1.9)^23^ was used for processing SAM and BAM files. Picard (v2.6.0) was used to remove duplicate reads. Peaks were called using Macs2 callpeak (v2.2.7.1)^24^ with default settings and bedtools intersect (v 2.31.1)^25^ used to exclude peaks that overlap with blacklisted regions (ENCODE). Motif analysis was performed using Homer (v4.11)^26^. Bigwig files were generated with deeptools (v 3.5.0)^27^ bamCoverage (-bl $blacklist --effectiveGenomeSize 2652783500 --binSize 10 -- smoothLength 50 --normalizeUsing CPM --extendReads 200) and visualised in IgV (v 2.14.1)^28^. A Bach1 union peak file was generated with bedtools merge and ChIPseeker (v 1.26.0) was used to annotate peaks for gene features^29^. Deeptools multiBigwigSummary was used to quantify the signal over the DNMT3A^R882H^ Bach1 peaks and ComplexHeatmap was used to integrate Bach1 ChIPseq signal and RNAseq data.

### Inductively coupled plasma mass spectrometry

OCI-AML3 cells were treated or not with SA (500μM) for 4 days. OCI-AML3-Cas9 sg Scr or sg2 ALAS1 cells were treated with doxycycline (1 μg/ml) to induce sgRNA expression for 7 days. For each sample, 1 x 10^7^ cells were harvested in 500 μL of cold HEPES (10 mM, pH 7.5) containing protease inhibitors cocktail without EDTA (Roche, USA) and phosphatase inhibitors cocktail (Roche, USA). Then, the cells were homogenized with 20 passages through a 25 G needle and centrifugation at 10,000 g for 10 min at 4°C. The supernatants were separated, and the protein concentration determined using the BCA Protein Assay Kit (Pierce, USA) and supernatants were aliquoted for Inductively coupled plasma mass spectrometry (ICP-MS) and stored at -80°C until use. 50 μL of sample (total homogenate or HEPES soluble fraction) was lyophilized for 12 h and digested with 50 µL of Nitric Acid (HNO_3_) (65% Suprapur, Merck) overnight at room temperature. Samples were further digested by heating at 90°C for 20 minutes using a heating block. The average reduced volume was determined, and the samples were further diluted with 1% HNO_3_ diluent. Measurements were made using an Agilent 8800 series ICP-MS instrument under routine multi-element operating conditions using a Helium Reaction Gas Cell. The instrument was calibrated using 0, 5, 10, 50, 100 and 500 ppb of certified multi-element ICP-MS standard calibration solutions (ICP-MS-CAL2-1, ICP-MS-CAL-3 and ICP-MS-CAL-4, AccuStandard) for a range of elements. Used a certified internal standard solution containing 200 ppb of Yttrium (Y89) as an internal control (ICP-MS-IS-MIX1-1, AccuStandard). Results are expressed as micrograms of copper per litre (μg/L).

### Immunofluorescence staining and confocal microscopy

OCI-AML3 cells transduced with ALAS1 or sg Scr sgRNAS and treated with doxycycline to induce sgRNA expression cells were seeded at a density of 1x10^5^ per ml onto sterile, poly-L-lysine coated (1mg/mL, P1274, Sigma) glass coverslips in a 24 well plate. Cells were allowed to attach and grow for 24 hours before fixation with 4% paraformaldehyde in PBS for 15min at room temperature. The coverslips were gently washed with PBS three times before incubation with 0.1% Triton X-100 in PBS for 15min at room temperature. The coverslips were gently washed again three times with PBS. The coverslips were blocked with 1% BSA in PBS for 1 hour at room temperature. Further washing with PBS before coverslips were transferred to be placed face down on parafilm with primary antibody diluted in 1% BSA in PBS overnight at 4°C (anti-ATP7a rabbit 1:400; gift from M. Greenough, Florey Institute, Australia). The coverslips were transferred back to their wells and gently washed three times with PBS before being transferred again to parafilm containing the secondary antibody diluted in 1% BSA in PBS (Alexa Fluor 594 goat anti rabbit IgG H+L 1:500, A11012, Thermo Fisher Scientific) for 2 hours at room temperature. The coverslips were transferred back to their wells and gently washed three times with PBS. DAPI (0.5ug/mL, D1306, Thermo Fisher Scientific) used to stain the nuclei was diluted in PBS and added to each well for 5 mins before being gently washed off. The coverslips were mounted onto a glass slide with Prolong Diamond Antifade Mounting Media (P36961, Thermo Fisher Scientific) and allowed to cure in the dark at room temperature. Images were acquired with a Leica SP8 DMI6000 inverted laser scanning microscope using a 63x/1.4NA oil objective lens. A Diode 405 laser was used to excite DAPI dye at 405nm, and emitted fluorescence was captured by a PMT detector. An Argon laser was used to excite the GFP protein. A DPSS 561 laser was used to excite the Alexa Fluor 594 dye at 561nm. Both the GFP and 594 emitted fluorescence were collected using a HyD detector.

### Quantification of total cellular heme using oxalic acid extraction

5x10^5^ (primary AML cells) or 5x10^6^ (AML cell lines and Hoxb8 cells) cells were aliquoted into tubes and centrifuged at 400 x g for 4 minutes. The supernatant was aspirated, and pellets were resuspended in 1 mL of PBS. The samples were centrifuged again at 400 x g for 4 minutes, the supernatant was aspirated, and pellets were resuspended in 100 µl of PBS. Standard samples were generated by serial dilution of hemin. 900µl of 2 M oxalic acid was added to each sample. The samples were divided into two 500µl aliquots. One aliquot of each sample was heated at 95°C for 30 minutes (boiled). The other aliquot was incubated at room temperature in darkness for 30 minutes (unboiled). The samples were allowed to cool at room temperature and centrifuged at 12000 RPM for 2 minutes. 150µl aliquots of each sample were analysed in technical duplicate on a Cytation 3 cell imaging multi-mode reader (BioTek) with excitation of 405nM and emission of 608nM. Total cellular heme content was calculated by subtracting the signal of unboiled samples from boiled samples and referenced to the hemin standard curve.

### 5-ALA flux assay

2 x 10^4^ cells were seeded in a 96-well plate and treated with 5-aminolevulinic acid (5-ALA) in technical duplicate for 3hrs at 37°C. For pulse chase experiments, cells were washed and incubated for an additional 16hrs. PpIX fluorescence was measured by flow cytometry with a 405nM laser and a 710/50 filter.

## Supplemental information

Table S1: RNA-seq of c-Kit+ DIN cells treated with doxycycline for 5 or 14 days.

Table S2: RNA-seq of HoxB8 EV and DNMT3A^R882H^ cells.

Table S3: All Bach1 CHIP-seq peaks in HoxB8 DNMT3A^R882H^ cells.

Table S4: GC-MS of HoxB8 EV and DNMT3A^R882H^ cells.

Table S5: RNA-seq of ALAS1 KO or Succinylacetone treatment in OCI-AML3 cells.

Table S6: MAGeCK RRA output of OCI-AML3 CRISPR screen under altered heme conditions.

Table S7: List of DNA sequences and constructs used in this study.

Table S8: Primary AML patient information.

Table S9: List of sgRNA sequences in metabolic focussed CRISPR library.

## References

1 Swenson, S. A. et al. From Synthesis to Utilization: The Ins and Outs of Mitochondrial Heme. Cells 9 (2020).

2 Dunaway, L. S., Loeb, S. A., Petrillo, S., Tolosano, E. & Isakson, B. E. Heme metabolism in non-erythroid cells. J Biol Chem 300, 107132 (2024).

3 Tsvetkov, P. et al. Copper induces cell death by targeting lipoylated TCA cycle proteins. Science 375, 1254–1261 (2022).

4 Hanahan, D. & Weinberg, R. A. Hallmarks of Cancer: The Next Generation. Cell 144, 646–674 (2011).

5 DeBerardinis, R. J., Lum, J. J., Hatzivassiliou, G. & Thompson, C. B. The biology of cancer: metabolic reprogramming fuels cell growth and proliferation. Cell Metab 7, 11–20 (2008).

6 Kinnaird, A., Zhao, S., Wellen, K. E. & Michelakis, E. D. Metabolic control of epigenetics in cancer. Nat Rev Cancer 16, 694–707 (2016).

7 Green, D. R., Galluzzi, L. & Kroemer, G. Metabolic control of cell death. Science 345, 1250256 (2014).

8 Stein, E. M. et al. Enasidenib in mutant IDH2 relapsed or refractory acute myeloid leukemia. Blood 130, 722–731 (2017).

9 DiNardo, C. D. et al. Durable Remissions with Ivosidenib in IDH1-Mutated Relapsed or Refractory AML. N Engl J Med 378, 2386–2398 (2018).

10 Pollyea, D. A. et al. Venetoclax with azacitidine disrupts energy metabolism and targets leukemia stem cells in patients with acute myeloid leukemia. Nat Med 24, 1859–1866 (2018).

11 Stine, Z. E., Schug, Z. T., Salvino, J. M. & Dang, C. V. Targeting cancer metabolism in the era of precision oncology. Nat Rev Drug Discov 21, 141–162 (2022).

12 Gu, Y. et al. IDH1 mutation contributes to myeloid dysplasia in mice by disturbing heme biosynthesis and erythropoiesis. Blood 137, 945–958 (2021).

13 Detraux, D. et al. A critical role for heme synthesis and succinate in the regulation of pluripotent states transitions. eLife 12 (2023).

14 Lin, K. H. et al. Systematic Dissection of the Metabolic-Apoptotic Interface in AML Reveals Heme Biosynthesis to Be a Regulator of Drug Sensitivity. Cell Metab. 29, 1217–1231.e1217 (2019).

15. Cancer Genome Atlas Research, N., et al. Genomic and epigenomic landscapes of adult de novo acute myeloid leukemiaN Engl J Med 368, 2059-2074 (2013).

16 Papaemmanuil, E. et al. Genomic Classification and Prognosis in Acute Myeloid Leukemia. N Engl J Med 374, 2209–2221 (2016).

17 Burd, A. et al. Precision medicine treatment in acute myeloid leukemia using prospective genomic profiling: feasibility and preliminary efficacy of the Beat AML Master Trial. Nat Med 26, 1852–1858 (2020).

18 Bhuva, D. D., Cursons, J. & Davis, M. J. Stable gene expression for normalisation and single-sample scoring. Nucleic Acids Res 48, e113 (2020).

19 Foroutan, M. et al. Single sample scoring of molecular phenotypes. BMC Bioinformatics 19, 404 (2018).

20 Raffel, S. et al. Quantitative proteomics reveals specific metabolic features of acute myeloid leukemia stem cells. Blood 136, 1507–1519 (2020).

21 Redecke, V. et al. Hematopoietic progenitor cell lines with myeloid and lymphoid potential. Nat Meth 10, 795–803 (2013).

22 Basilico, S. et al. Dissecting the early steps of MLL induced leukaemogenic transformation using a mouse model of AML. Nat Commun 11, 1407–1415 (2020).

23 Xu, J. J. et al. Genome-wide screening identifies cell-cycle control as a synthetic lethal pathway with SRSF2P95H mutation. Blood Adv 6, 2092–2106 (2022).

24 Challen, G. A. et al. Dnmt3a is essential for hematopoietic stem cell differentiation. Nat Genet 44, 23–31 (2011).

25 Hanna, D. A. et al. Heme dynamics and trafficking factors revealed by genetically encoded fluorescent heme sensors. Proc Natl Acad Sci U S A 113, 7539–7544 (2016).

26 Ogawa, K. et al. Heme mediates derepression of Maf recognition element through direct binding to transcription repressor Bach1. EMBO J 20, 2835–2843 (2001).

27 Kaasik, K. & Lee, C. C. Reciprocal regulation of haem biosynthesis and the circadian clock in mammals. Nature 430, 467–471 (2004).

28 Raghuram, S. et al. Identification of heme as the ligand for the orphan nuclear receptors REV-ERBalpha and REV-ERBbeta. Nat Struct Mol Biol 14, 1207–1213 (2007).

29 Gray, L. T. et al. G-quadruplexes Sequester Free Heme in Living Cells. Cell Chem Biol 26, 1681–1691 e1685 (2019).

30 Puram, R. V. et al. Core Circadian Clock Genes Regulate Leukemia Stem Cells in AML. Cell 165, 303–316 (2016).

31 Zenke-Kawasaki, Y. et al. Heme induces ubiquitination and degradation of the transcription factor Bach1. Mol Cell Biol 27, 6962–6971 (2007).

32 Igarashi, K., Nishizawa, H., Saiki, Y. & Matsumoto, M. The transcription factor BACH1 at the crossroads of cancer biology: From epithelial-mesenchymal transition to ferroptosis. J Biol Chem 297, 101032 (2021).

33 Hao, S. et al. Chronic intermittent hypoxia promoted lung cancer stem cell-like properties via enhancing Bach1 expression. Respir Res 22, 58–12 (2021).

34 Fujihara, K. M. et al. Eprenetapopt triggers ferroptosis, inhibits NFS1 cysteine desulfurase, and synergizes with serine and glycine dietary restriction. Sci Adv 8, eabm9427 (2022).

35 Dixon, S. J. et al. Ferroptosis: an iron-dependent form of nonapoptotic cell death. Cell 149, 1060–1072 (2012).

36 Barbie, D. A. et al. Systematic RNA interference reveals that oncogenic KRAS-driven cancers require TBK1. Nature 462, 108–112 (2009).

37 Freeman, S. L. et al. Heme binding to human CLOCK affects interactions with the E-box. Proc Natl Acad Sci U S A 116, 19911–19916 (2019).

38 Guzman, M. L. et al. Nuclear factor-kappaB is constitutively activated in primitive human acute myelogenous leukemia cells. Blood 98, 2301–2307 (2001).

39 Kagoya, Y. et al. Positive feedback between NF-kappaB and TNF-alpha promotes leukemia-initiating cell capacity. J Clin Invest 124, 528–542 (2014).

40 Ebert, P. S., Hess, R. A., Frykholm, B. C. & Tschudy, D. P. Succinylacetone, a potent inhibitor of heme biosynthesis: effect on cell growth, heme content and delta-aminolevulinic acid dehydratase activity of malignant murine erythroleukemia cells. Biochem Biophys Res Commun 88, 1382–1390 (1979).

41 Wang, Z. et al. Methionine is a metabolic dependency of tumor-initiating cells. Nat Med 25, 825–837 (2019).

42 Chen, C.-C. et al. Vitamin B6 Addiction in Acute Myeloid Leukemia. Cancer Cell 37, 71–83.e78 (2020).

43 Wei, Y. et al. SLC5A3-Dependent Myo-inositol Auxotrophy in Acute Myeloid Leukemia. Cancer Discov 12, 450–467 (2022).

44 Gamble, J. T., Dailey, H. A. & Marks, G. S. N-Methylprotoporphyrin is a more potent inhibitor of recombinant human than of recombinant chicken ferrochelatase. Drug Metab Dispos 28, 373–375 (2000).

45 Drexler, H. G., Gaedicke, G. & Minowada, J. Erythroleukemia cell lines HEL and K-562: changes in isoenzyme profiles and morphology during induction of differentiation. Hematol Oncol 4, 163–174 (1986).

46 Liu, X. et al. Actin cytoskeleton vulnerability to disulfide stress mediates disulfidptosis. Nat Cell Biol 25, 404–414 (2023).

47 Kurtoglu, M. et al. Under normoxia, 2-deoxy-D-glucose elicits cell death in select tumor types not by inhibition of glycolysis but by interfering with N-linked glycosylation. Mol Cancer Ther 6, 3049–3058 (2007).

48 Maddocks, O. D. et al. Serine starvation induces stress and p53-dependent metabolic remodelling in cancer cells. Nature 493, 542–546 (2013).

49 Li, W. et al. Quality control, modeling, and visualization of CRISPR screens with MAGeCK-VISPR. Genome Biol 16, 281 (2015).

50 Li, W. et al. MAGeCK enables robust identification of essential genes from genome-scale CRISPR/Cas9 knockout screens. Genome Biol 15, 819–812 (2014).

51 Rowland, E. A., Snowden, C. K. & Cristea, I. M. Protein lipoylation: an evolutionarily conserved metabolic regulator of health and disease. Curr Opin Chem Biol 42, 76–85 (2018).

52 Dreishpoon, M. B. et al. FDX1 regulates cellular protein lipoylation through direct binding to LIAS. J Biol Chem, 105046 (2023).

53 Zulkifli, M., Okonkwo, A. U. & Gohil, V. M. FDX1 Is Required for the Biogenesis of Mitochondrial Cytochrome c Oxidase in Mammalian Cells. J Mol Biol 435, 168317 (2023).

54 Bertinato, J., Sherrard, L. & Plouffe, L. J. Decreased erythrocyte CCS content is a biomarker of copper overload in rats. Int J Mol Sci 11, 2624–2635 (2010).

55 Bertinato, J. & L’Abbe, M. R. Copper modulates the degradation of copper chaperone for Cu,Zn superoxide dismutase by the 26 S proteosome. J Biol Chem 278, 35071–35078 (2003).

56 Xie, L. & Collins, J. F. Copper stabilizes the Menkes copper-transporting ATPase (Atp7a) protein expressed in rat intestinal epithelial cells. Am J Physiol Cell Physiol 304, C257–C262 (2013).

57 Birsoy, K. et al. An Essential Role of the Mitochondrial Electron Transport Chain in Cell Proliferation Is to Enable Aspartate Synthesis. Cell 162, 540–551 (2015).

58 Baccelli, I. et al. Mubritinib Targets the Electron Transport Chain Complex I and Reveals the Landscape of OXPHOS Dependency in Acute Myeloid Leukemia. Cancer Cell 36, 84–99.e88 (2019).

59 DeWaal, D. et al. Hexokinase-2 depletion inhibits glycolysis and induces oxidative phosphorylation in hepatocellular carcinoma and sensitizes to metformin. Nat Commun 9, 446 (2018).

60 Rodriguez-Zabala, M. et al. Combined GLUT1 and OXPHOS inhibition eliminates acute myeloid leukemia cells by restraining their metabolic plasticity. Blood Adv 7, 5382–5395 (2023).

61 Siebeneicher, H. et al. Identification and Optimization of the First Highly Selective GLUT1 Inhibitor BAY-876. ChemMedChem 11, 2261–2271 (2016).

62 Lagadinou, E. D. et al. BCL-2 inhibition targets oxidative phosphorylation and selectively eradicates quiescent human leukemia stem cells. Cell Stem Cell 12, 329–341 (2013).

63 Moison, C. et al. SF3B1 mutations provide genetic vulnerability to copper ionophores in human acute myeloid leukemia. Sci Adv 10, eadl4018 (2024).

64 Solier, S. et al. A druggable copper-signalling pathway that drives inflammation. Nature, 1–9 (2023).

65 Kao, Y. R. et al. An iron rheostat controls hematopoietic stem cell fate. Cell Stem Cell (2024).

66 Sallman, D. A. et al. Eprenetapopt (APR-246) and Azacitidine in TP53-Mutant Myelodysplastic Syndromes. J Clin Oncol 39, 1584–1594 (2021).

67 Zhang, H. et al. Susceptibility of acute myeloid leukemia cells to ferroptosis and evasion strategies. Front Mol Biosci 10, 1275774 (2023).

## References

1 Redecke, V. et al. Hematopoietic progenitor cell lines with myeloid and lymphoid potential. Nat Meth 10, 795–803 (2013).

2 So, J. et al. Inhibition of pyrimidine biosynthesis targets protein translation in acute myeloid leukemia. EMBO Mol Med 14, e15203 (2022).

3 Zuber, J. et al. An integrated approach to dissecting oncogene addiction implicates a Myb-coordinated self-renewal program as essential for leukemia maintenance. Genes Dev. 25, 1628–1640 (2011).

4 Gruber, E. et al. Inhibition of mutant IDH1 promotes cycling of acute myeloid leukemia stem cells. Cell Rep 40, 111182 (2022).

5 Birsoy, K. et al. An Essential Role of the Mitochondrial Electron Transport Chain in Cell Proliferation Is to Enable Aspartate Synthesis. Cell 162, 540–551 (2015).

6 Corcoran, C. C., Grady, C. R., Pisitkun, T., Parulekar, J. & Knepper, M. A. From 20th century metabolic wall charts to 21st century systems biology: database of mammalian metabolic enzymes. Am J Physiol Renal Physiol 312, F533–F542 (2017).

7 Doench, J. G. et al. Optimized sgRNA design to maximize activity and minimize off-target effects of CRISPR-Cas9. Nat Biotech 34, 184–191 (2016).

8 Martin, M. Cutadapt removes adapter sequences from high-throughput sequencing reads. EMBnet.journal 17, 10–12 (2011).

9 Li, W. et al. MAGeCK enables robust identification of essential genes from genome-scale CRISPR/Cas9 knockout screens. Genome Biol 15, 819–812 (2014).

10 Kim, D., Langmead, B. & Salzberg, S. L. HISAT: a fast spliced aligner with low memory requirements. Nat Meth 12, 357–360 (2015).

11 Liao, Y., Smyth, G. K. & Shi, W. featureCounts: an efficient general purpose program for assigning sequence reads to genomic features. Bioinformatics 30, 923–930 (2014).

12 McCarthy, D. J., Chen, Y. & Smyth, G. K. Differential expression analysis of multifactor RNA-Seq experiments with respect to biological variation. Nucleic Acids Res 40, 4288–4297 (2012).

13 Ritchie, M. E. et al. limma powers differential expression analyses for RNA-sequencing and microarray studies. Nucleic Acids Res 43, e47 (2015).

81. Korotkevich, G., et al. Fast gene set enrichment analysis. bioRxiv (2021).

15 Hoadley, K. A. et al. Cell-of-Origin Patterns Dominate the Molecular Classification of 10,000 Tumors from 33 Types of Cancer. Cell 173, 291–304 e296 (2018).

16 Tyner, J. W. et al. Functional genomic landscape of acute myeloid leukaemia. Nature, s**562**, 526–531 (2018).

17 Hansen, K. D., Irizarry, R. A. & Wu, Z. Removing technical variability in RNA-seq data using conditional quantile normalization. Biostatistics 13, 204–216 (2012).

18 Foroutan, M. et al. Single sample scoring of molecular phenotypes. BMC Bioinformatics 19, 404 (2018).

19 Bhuva, D. D., Cursons, J. & Davis, M. J. Stable gene expression for normalisation and single-sample scoring. Nucleic Acids Res 48, e113 (2020).

20 Gu, Z., Eils, R. & Schlesner, M. Complex heatmaps reveal patterns and correlations in multidimensional genomic data. Bioinformatics 32, 2847–2849 (2016).

21 Langmead, B. & Salzberg, S. L. Fast gapped-read alignment with Bowtie 2. Nat Methods 9, 357–359 (2012).

22 Langmead, B., Wilks, C., Antonescu, V. & Charles, R. Scaling read aligners to hundreds of threads on general-purpose processors. Bioinformatics 35, 421–432 (2019).

23 Li, H. et al. The Sequence Alignment/Map format and SAMtools. Bioinformatics 25, 2078–2079 (2009).

24 Zhang, Y. et al. Model-based analysis of ChIP-Seq (MACS). Genome Biol 9, R137 (2008).

25 Quinlan, A. R. & Hall, I. M. BEDTools: a flexible suite of utilities for comparing genomic features. Bioinformatics 26, 841–842 (2010).

26 Heinz, S. et al. Simple combinations of lineage-determining transcription factors prime cis-regulatory elements required for macrophage and B cell identities. Mol Cell 38, 576–589 (2010).

27 Ramirez, F., Dundar, F., Diehl, S., Gruning, B. A. & Manke, T. deepTools: a flexible platform for exploring deep-sequencing data. Nucleic Acids Res 42, W187–191 (2014).

28 Robinson, J. T. et al. Integrative genomics viewer. Nat Biotechnol 29, 24–26 (2011).

29 Yu, G., Wang, L. G. & He, Q. Y. ChIPseeker: an R/Bioconductor package for ChIP peak annotation, comparison and visualization. Bioinformatics 31, 2382–2383 (2015).

