## Extended figures for "Inhibition of heme biosynthesis triggers cuproptosis in acute myeloid leukaemia"

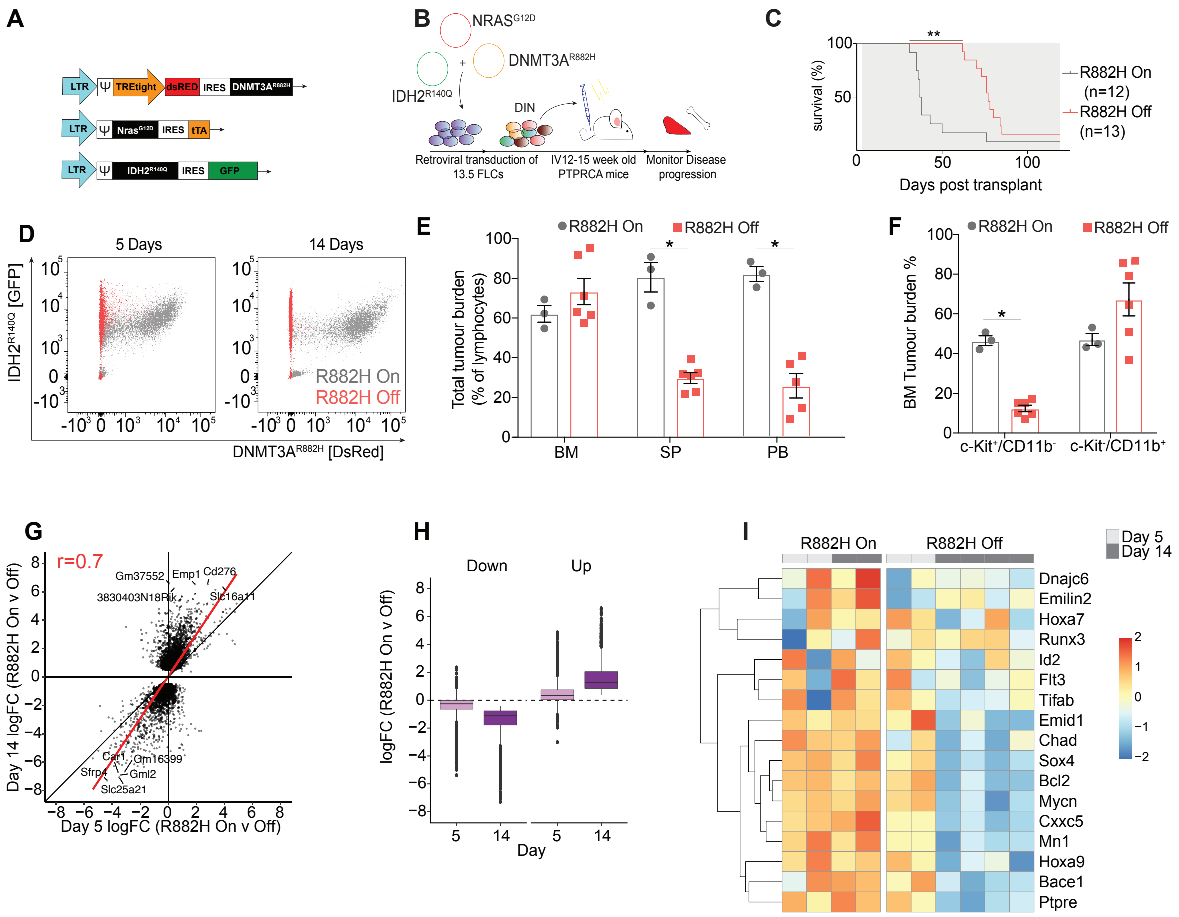


***Extended Data Figure 1. Characterisation of the DIN mouse model.***

**A.** Schematic of retroviral constructs used to generate the multi-allelic DIN AML model. DNMT3AR882H expression is doxycycline inducible; IDH2$^{R140Q}$ and NRAS$^{G12D}$ expression are constitutive. **B-C.** Ptprca recipient mice were engrafted with cells transduced with DNMT3A$^{R882H}$. **C.** Kaplan-Meier survival curve. Grey denotes when mice were administered normal or doxycyline chow and water. **E-F.** FACS analysis of Ptprca recipient mice engrafted with DIN leukaemias and treated or not with doxycycline for 5 or 14 days (n = 3-6 mice). Significance was assessed with unpaired t-test. * $p < 0.05$. BM, bone marrow. SP, spleen. PB, peripheral blood. **G-I.** RNAseq analysis of cKit+CD11b- DIN progenitors from tumour bearing mice untreated or treated with doxycycline for 5 or 14 days (n = 2-4 mice/group). **G.** Correlation of gene expression changes of DEGs (FDR < 0.05 & |logFC| > 0.5 at either timepoint). **H.** Magnitude of gene expression changes from G. **I.** Heatmap showing expression of canonical DNMT3A$^{R882H}$ target genes.

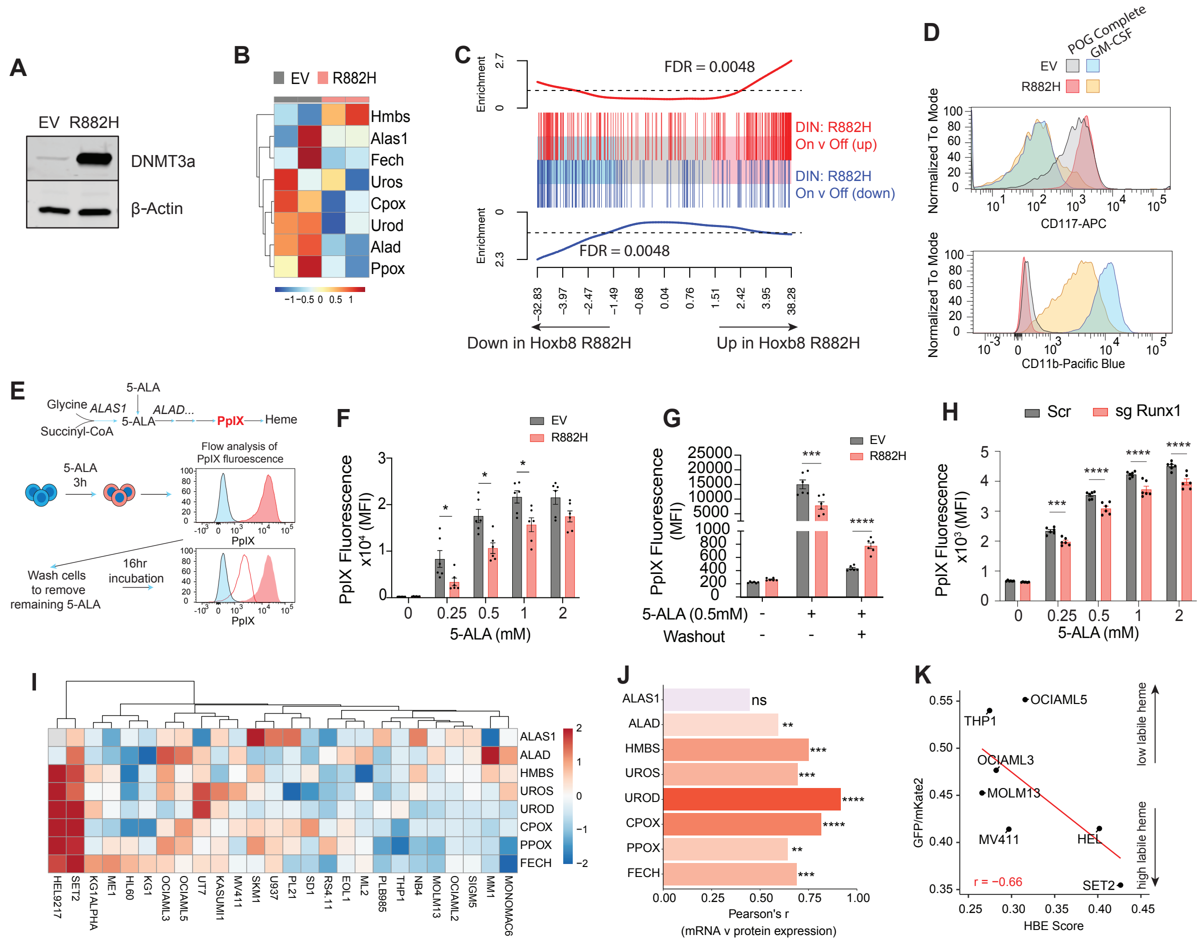


***Extended Data Figure 2. AML driver mutations reduce heme levels.***

**A-D, F-G.** Hoxb8 cells transduced with empty vector (control) or DNMT3$^{AR882H}$. **A.** Western blot for the indicated proteins. **B-C.** Gene expression analysed by RNAseq. **B.** Heatmap of HBE expression in Hoxb8 cells. **C.** Barcode plot showing enrichment of transcriptional signatures derived from cKit+CD11b- DIN progenitors (Day 14 R882H On v Off DEGs: FDR < 0.01 & |logFC|>2) in Hoxb8 DNMT3A$^{R882H}$ cells. Significance determined by ROAST. **D.** Cells were cultured in complete media or with GM-CSF to promote differentiation. Expression of cKit and CD11b were analysed by FACS. **E.** Schematic for 5-ALA FACS assay to measure flux through the heme biosynthesis pathway. De novo heme production commences through the condensation of succinyl-CoA and glycine into 5-ALA. Heme pathway activity was determined by the capacity of cells to convert exogenous 5-ALA into the fluorescent intermediate, protoporphyrin IX (PpIX) (Pulse). 5-ALA treated cells were washed to remove excess 5-ALA and measure their capacity to remove the remaining PpIX (Chase). **F-H.** 5-ALA assay to quantify PpIX fluorescence by FACS. n = 3 biological replicates. Error bars represent mean ± SEM. Significance was assessed with a multiple comparison adjusted t-test; * $p < 0.05$, ** $p < 0.01$, *** $p < 0.001$, **** $p < 0.0001$. **F.** Hoxb8 cells were incubated with increasing concentrations of 5-ALA and PpIX fluorescence was quantified by FACS. **G.** Hoxb8 cells were incubated with 5-ALA (0.5 mM) for 3 hours (pulse), 5-ALA was washed out, cells were cultured for 16 hours (chase), and PpIX fluorescence was quantified by FACS. **H.** Cas9 expressing Hoxb8 cells transduced with scrambled or a Runx1 targeting sgRNA were incubated with increasing concentrations of 5-ALA and PpIX fluorescence was quantified by FACS. **I.** Heatmap of protein levels of HBE genes in AML cell lines[1]. **J.** Pearson's correlation coefficient r and corresponding p-values (denoted as stars) when comparing RNA (CCLE) and protein levels of HBE genes in AML cell lines1. * $p < 0.05$, ** $p < 0.01$, *** $p < 0.001$, **** $p < 0.0001$. **K.** Correlation plot between HBE score and labile heme in AML cell lines. Increased GFP/mKate ratio indicates lower labile heme levels[2]. r represents Pearson correlation coefficient.

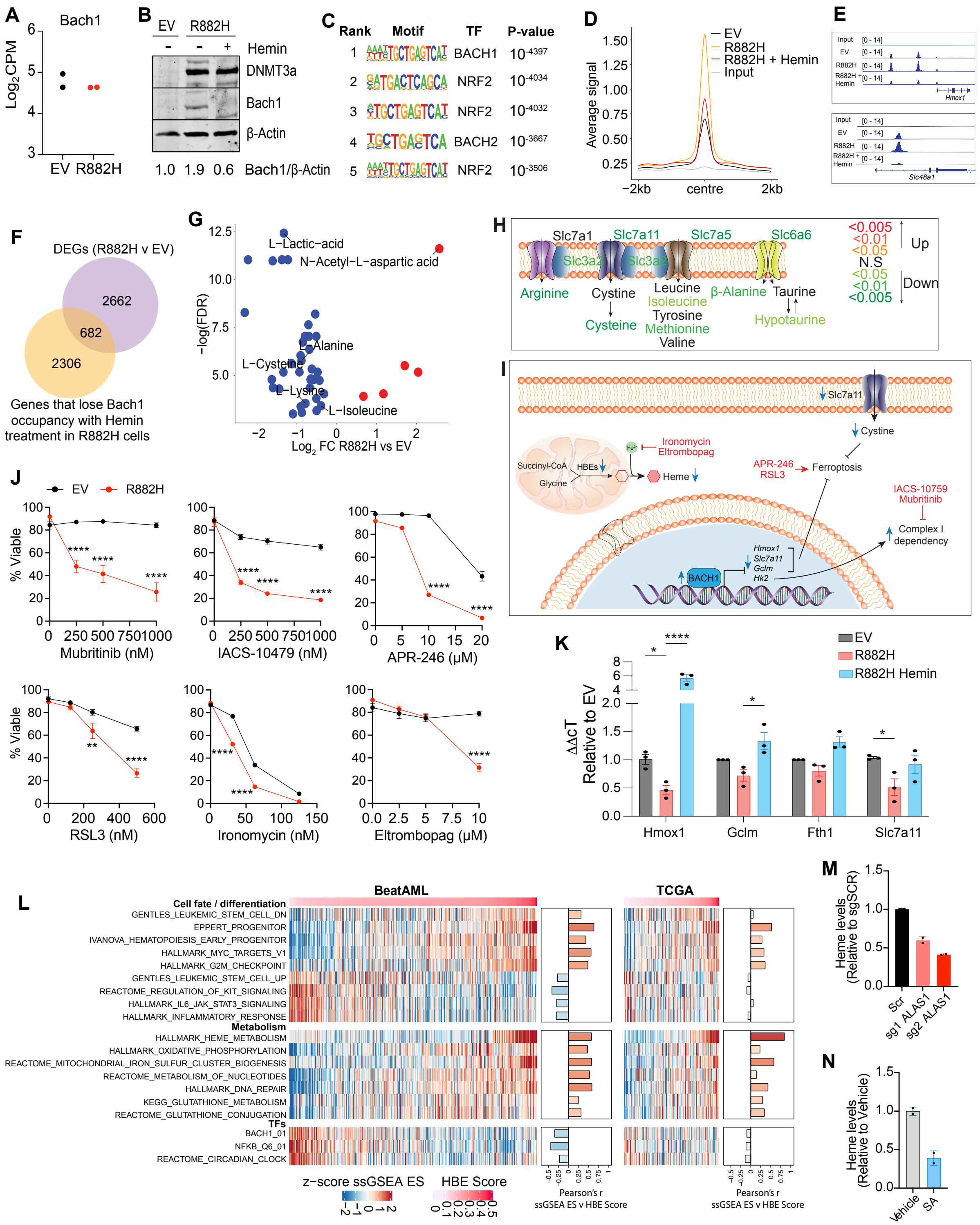


***Extended Data Figure 3. The heme/bach1 axis regulates metabolic and self-renewal gene expression programs.***

**A.** BACH1 RNA quantified by RNAseq in Hoxb8 cells. (n = 2 biological replicates). **B.** Hoxb8 cells transduced with empty vector (control) or DNMT3A$^{R882H}$ and treated with or without hemin (40 µM). **C-E.** ChIPseq for Bach1 in Hoxb8 cells. **C.** HOMER analysis of transcription factor motifs enriched within all Bach1 ChIPseq peaks. **D.** Bach1 ChIPseq signal centred on all Bach1 peaks in DNMT3A$^{R882H}$ transduced cells treated with vehicle. **E.** IGV tracks of canonical Bach1 target genes, Hmox1 and Slc48a1. **F.** Venn diagram of the overlap between heme-responsive Bach1 bound genes and DEGs (DNMT3A$^{R882H}$ v EV). **G.** Volcano plot showing differentially abundant metabolites (FDR $< 0.05$) quantified by GC-MS. **H.** Summary schematic of amino acids and protein transporters differentially regulated in Hoxb8 cells transduced with empty vector (control) or DNMT3A$^{R882H}$. Amino acid levels were quantified by GC-MS ($n = 5$ biological replicates; mean ± SEM; significance was assessed with a multiple comparison adjusted t-test). Transporter expression was quantified by RNAseq ($n = 2$ biological replicates). **I.** Reductions in heme levels results in Bach1 stabilisation and repression of Bach1 target genes which increases susceptibility of low heme cells to specific compounds (Red) by targeting processes such as iron and glutathione metabolism and Complex I. Blue arrows indicate the consequence of low cellular heme levels. Red arrows indicate pathways that are activated or inhibited by the indicated compound. **J.** Viability of Hoxb8 cells transduced with empty vector (EV) or DNMT3A$^{R882H}$ treated with mubritinib (72 hours), IACS-10759 (72 hours), APR-246 (24 hours), RSL3 (24 hours), ironomycin (24 hours) or eltrombopag (72 hours). $n = 3$ biological replicates. Error bars represent mean ± SEM. Significance was assessed with a multiple comparison adjusted t-test; * $p < 0.05$, ** $p < 0.01$, *** $p < 0.001$, **** $p < 0.0001$. **K.** qPCR analysis of selected genes containing Bach1 peaks. $n = 3$ biological replicates. Error bars represent mean ± SEM. Significance was assessed with a multiple comparison adjusted t-test * $p <0.05$, **** $p <0.0001$. **L.** ssGSEA of all AMLs from the BeatAML and TCGA datasets. **M-N.** Total heme levels in OCI-AML3 cells **M.** transduced with ALAS1 or scrambled sgRNAs and treated with doxycycline to induce sgRNA expression or **N)** SA for 7 days. $n = 2$ biological replicates. Error bars represent mean ± SD.

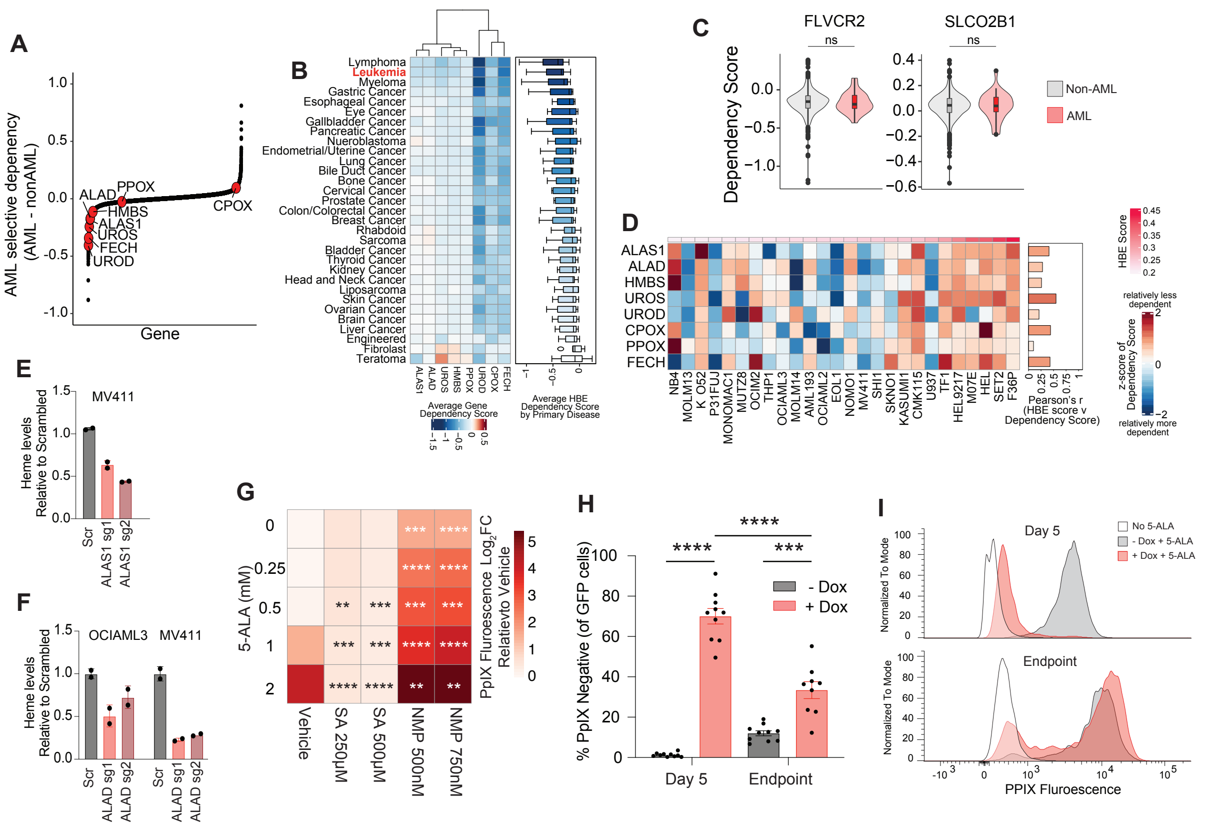


***Extended Data Figure 4: De novo heme biosynthesis is a selective leukaemia dependency.*** **A-D.** Analysis of DepMap data. **A.** AML selective dependencies, highlighting HBE genes. Lower score indicates greater selective dependency in AML. **B.** Dependency scores for HBE genes in cell lines from different cancer types. **C.** Dependency scores for genes encoding heme importers (FLVCR2, SLCO2B1) comparing AML and non-AML cell lines. Significance was assessed with a one-sided t-test. ns; $p \geq 0.05$. **D.** Heatmap depicting z-score of dependency scores of AML cell lines (blue indicates relatively more dependent). Pearson's r showing positive correlation of HBE expression (HBE score) and dependency (DepMap dependency score). **E-F.** Total heme levels in OCI-AML3 and MV4-11 cells transduced with ALAS1, ALAD or scrambled sgRNAs and treated with doxycycline to induce sgRNA expression. n = 2 biological replicates. Error bars represent mean +/- SD. **G.** OCI-AML3 were pretreated with SA or NMP for 72 hours and then incubated with 5-ALA for 3 hours and PpIX fluorescence quantified by FACS (n = 3 biological replicates). **H-I.** Ex vivo analysis of PpIX fluorescence. MV4-11 cells transduced with a doxycycline inducible ALAD sgRNA were harvested from mice after 5 days of doxycycline chow and water administration, or at ethical endpoint, then treated ex vivo with 5-ALA for 3 hours. n = 9-10 biological replicates. Error bars represent mean +/- SEM. **I.** Representative FACS plots after 5 days of doxycycline treatment or at disease endpoint.

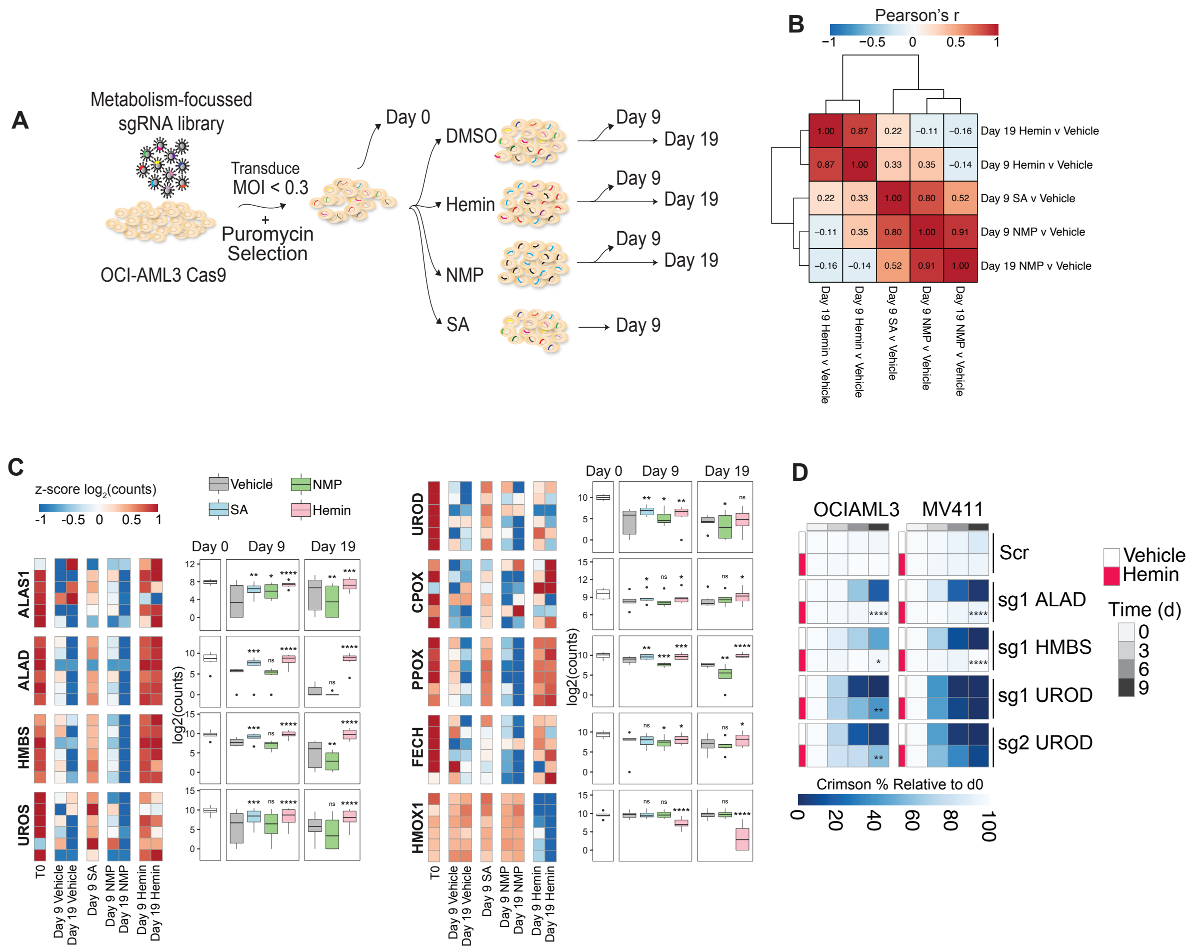


***Extended Data Figure 5. Manipulating cellular heme levels induces specific metabolic vulnerabilities.***

**A.** Schematic of CRISPR screen. **B.** Pearson's r denoting the correlation of the logFC of knockout hits (FDR < 0.05 in either drug condition relative to vehicle) between the different treatment conditions and timepoints. **C.** Heatmap and boxplot of normalised sgRNA counts of HBE genes and HMOX1 under Vehicle, SA, NMP and Hemin treatment conditions. Significance assessed using MAGECK-RRA p-values relative to vehicle. * $p < 0.05$, ** $p < 0.01$, *** $p < 0.001$, **** $p < 0.0001$. **D.** Competition assays of OCI-AML3 and MV4-11 cells transduced with sgRNAs targeting ALAD, HMBS or UROD and treated with Hemin. n = 2 biological replicates. Significance assessed by two-way ANOVA. * $p < 0.05$, ** $p < 0.01$, **** $p < 0.0001$.

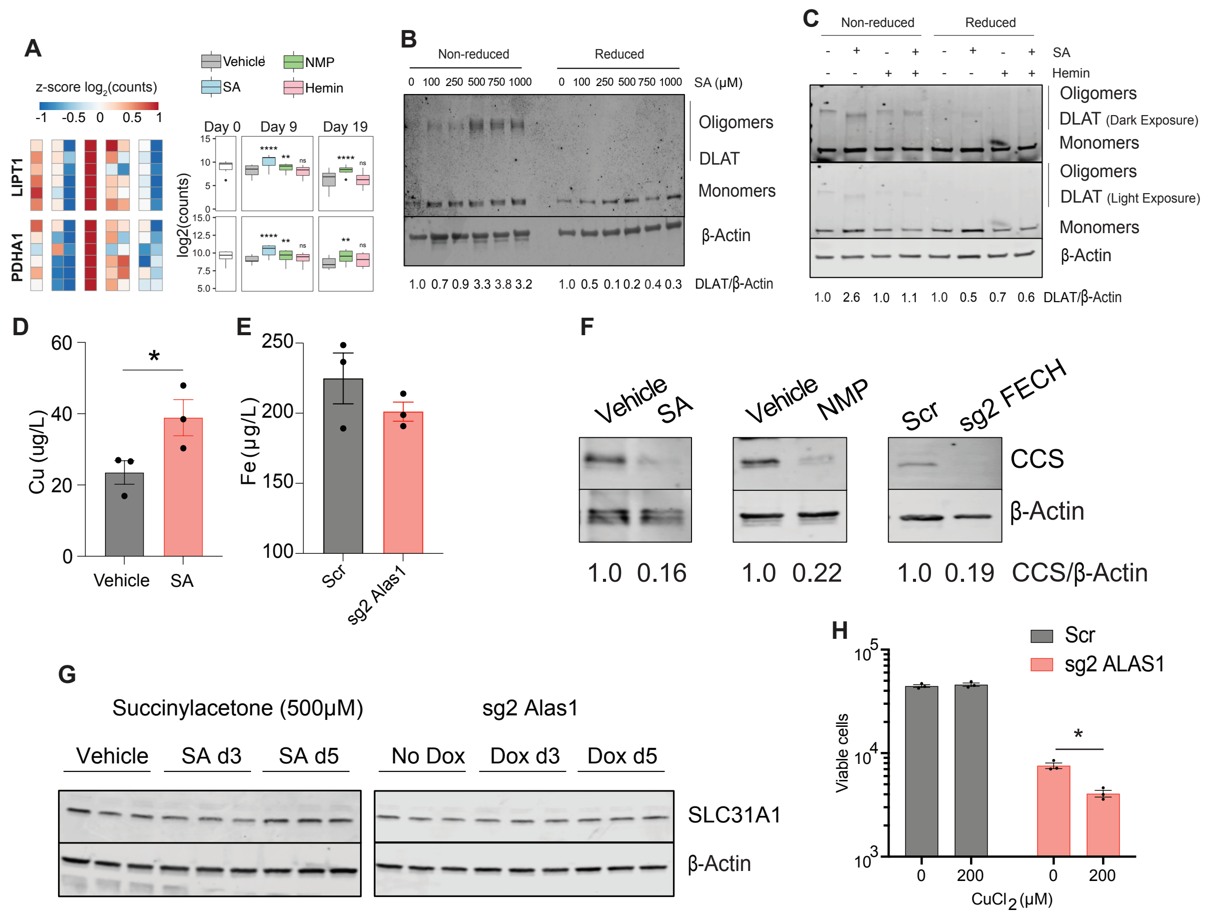


***Extended Data Figure 6: Heme starvation induces cuproptosis***

**A.** Heatmap and boxplot of normalised sgRNA counts of lipoic acid genes under Vehicle, SA, NMP and Hemin treatment conditions. Significance assessed using MAGECK-RRA p-values compared to vehicle. * p <0.05, ** p <0.01, *** p <0.001, **** p <0.0001. **B-C.** Western blot analysis OCI-AML3 cells treated with SA or vehicle and **C.** with or without hemin. **D-E.** Inductively coupled plasma mass spectrometry (ICP-MS) of **D.** copper levels in OCI-AML3 cells treated with SA or vehicle for 4 days or **E.** iron levels in OCI-AML3 cells expressing ALAS1 or scrambled sgRNAs for 7 days. n = 3 biological replicates. Error bars represent mean ± SEM. Significance was assessed using unpaired students T-test * p < 0.05, ** p < 0.01. **F.** Western blot of copper chaperone CCS in OCI-AML3 cells treated with SA, NMP or doxycycline to target FECH. **G.** Western blot analysis of copper importer SLC31A1 in OCI-AML3 cells transduced with inducible sgRNA targeting ALAS1 3, 5 days post doxycycline treatment or SA treatment (500 μM). **H.** Viability of OCI-AML3 cells expressing doxycycline-inducible ALAS1 or scrambled sgRNAs treated with copper chloride for 10 days. n = 3 biological replicates. Error bars represent mean ± SEM. Significance was assessed using by an unpaired students T-test * p < 0.05.

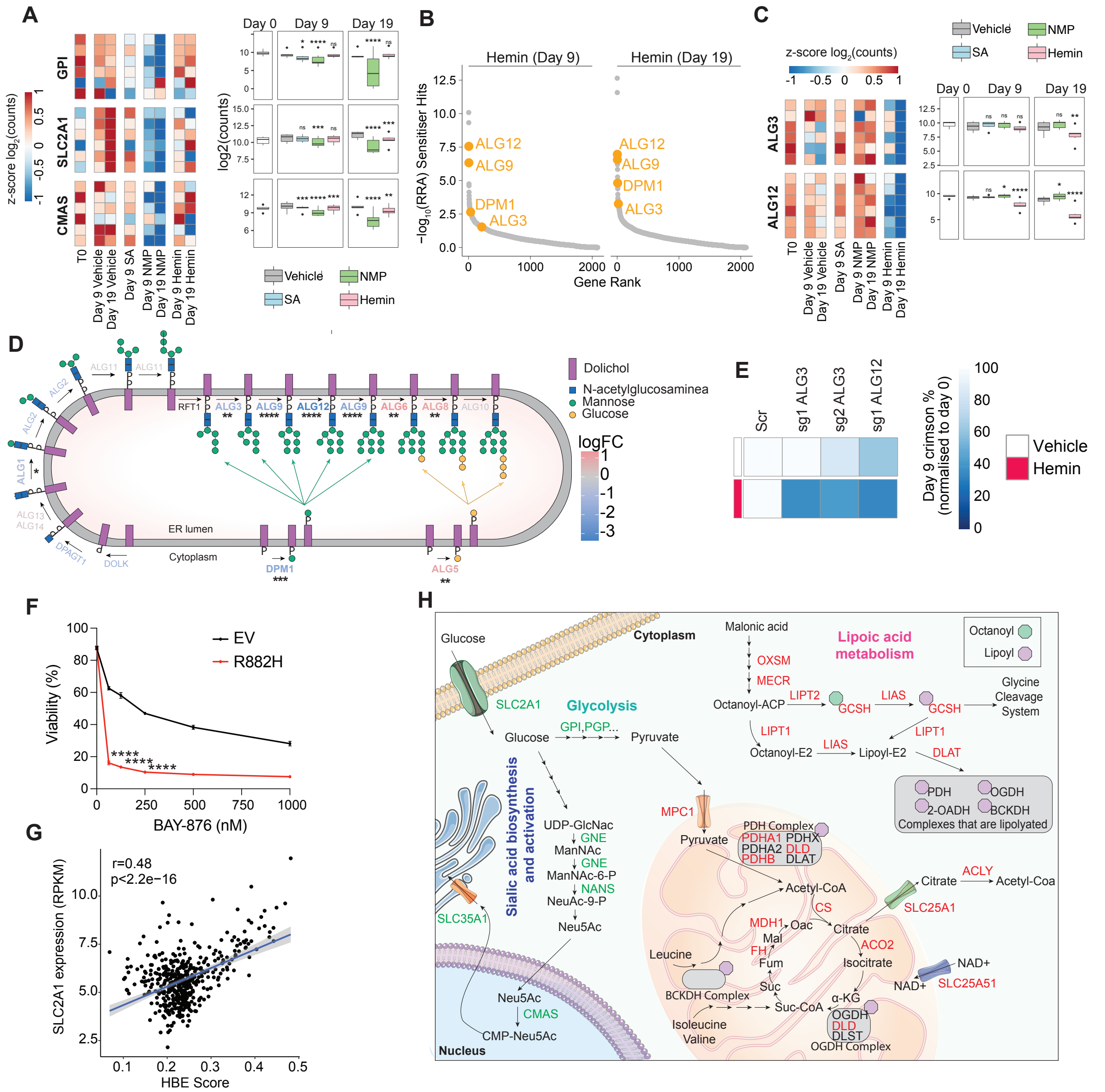


***Extended Data Figure 7. Cellular heme levels affect metabolic dependencies in AML.***

**A.** Heatmap and boxplot of normalised sgRNA counts of glycolysis and sialic acid genes under Vehicle, SA, NMP and Hemin treatment conditions. Significance assessed using MAGECK-RRA p-values compared to vehicle; * p <0.05, ** p <0.01, *** p <0.001, **** p <0.0001. **B.** MAGECK-RRA analysis identifying genes that increase the sensitivity of AML cells to heme supplementation by hemin. **C.** Heatmap and boxplot of normalised sgRNA counts of N-linked glycan genes under Vehicle, SA, NMP and Hemin treatment conditions. Significance assessed using MAGECK-RRA p-values compared to vehicle; * p <0.05, ** p <0.01, *** p <0.001, **** p <0.0001. **D.** Schematic summary of genetic hits associated with the N-linked glycan pathway. **E.** Proliferative competition assays of AML cells transduced with sgRNAs targeting ALG3 or ALG12 and treated with Hemin. n = 2 biological replicates. Significance was assessed by two-way ANOVA. **F.** Viability of Hoxb8 cells transduced with empty vector (EV) or DNMT3A$^{R882H}$ treated with BAY-876 (48 hours). n = 3 biological replicates. Error bars represent mean ± SEM. Significance was assessed with a multiple comparison adjusted t-test; **** $p < 0.0001$. **G.** Correlation plot of SLC2A1 expression and HBE score in the BEAT AML cohort. r represents Pearson correlation coefficient. **H.** Schematic summary of genetic hits associated with heme starvation. Genes highlighted in red and green are associated with resistance and sensitisation, respectively, under low heme conditions induced by SA or NMP.

### References


1 Jayavelu, A. K. *et al.* The proteogenomic subtypes of acute myeloid leukemia. *Cancer Cell* **40**, 301-317 (2022).

2 Hanna, D. A. *et al.* Heme dynamics and trafficking factors revealed by genetically encoded fluorescent heme sensors. *Proc Natl Acad Sci U S A* **113**, 7539-7544 (2016).
